# The L-lactate dehydrogenases of *Pseudomonas aeruginosa* are conditionally regulated but both contribute to survival during macrophage infection

**DOI:** 10.1101/2024.03.21.586142

**Authors:** Lindsey C. Florek, Xi Lin, Yu-Cheng Lin, Min-Han Lin, Arijit Chakraborty, Alexa Price-Whelan, Liang Tong, Laurence Rahme, Lars E.P. Dietrich

## Abstract

*Pseudomonas aeruginosa* is an opportunistic pathogen that thrives in environments associated with human activity, including soil and water altered by agriculture or pollution. Because L-lactate is a significant product of plant and animal metabolism, it is available to serve as a carbon source for *P. aeruginosa* in the diverse settings it inhabits. Here, we evaluate *P. aeruginosa*’s production and use of its redundant L-lactate dehydrogenases, termed LldD and LldA. We confirm that the protein LldR represses *lldD* and identify a new transcription factor, called LldS, that activates *lldA*; these distinct regulators and the genomic contexts of *lldD* and *lldA* contribute to their differential expression. We demonstrate that the *lldD* and *lldA* genes are conditionally controlled in response to lactate isomers as well as to glycolate and ◻-hydroxybutyrate, which, like lactate, are ◻-hydroxycarboxylates. We also show that *lldA* is induced when iron availability is low. Our examination of *lldD* and *lldA* expression across depth in biofilms indicates a complex pattern that is consistent with the effects of glycolate production, iron availability, and cross-regulation on enzyme preference. Finally, macrophage infection assays revealed that both *lldD* and *lldA* contribute to persistence within host cells, underscoring the potential role of L-lactate as a carbon source during *P. aeruginosa*-eukaryote interactions. Together, these findings help us understand the metabolism of a key resource that may promote *P. aeruginosa*’s success as a resident of contaminated environments and animal hosts.

**Importance:** *Pseudomonas aeruginosa* is a major cause of lung infections in people with cystic fibrosis, hospital-acquired infections, and wound infections. It consumes L-lactate, which is found at substantial levels in human blood and tissues. In this study, we investigated the spatial regulation of two redundant enzymes, called LldD and LldA, which enable L-lactate metabolism in *P. aeruginosa* biofilms. We uncovered mechanisms and identified compounds that control *P. aeruginosa*’s LldD/LldA preference. We also showed that both enzymes contribute to its ability to survive within macrophages, a behavior that is thought to augment the chronicity and recalcitrance of infections. Our findings shed light on a key metabolic strategy used by *P. aeruginosa* and have the potential to inform the development of therapies targeting bacterial metabolism during infection.

## Introduction

Lactate is a small organic compound and a metabolite that is present in diverse environments. In the rhizosphere or within animal hosts, lactate can constitute a major carbon source for commensal and pathogenic microbes (1–4). The bacterial “NAD-independent” lactate dehydrogenases, referred to here as “iLDH”s, enable growth on lactate by oxidizing it to pyruvate (5). iLDHs are specific for either the D-or L-isomer of lactate and bacteria generally show variation in their complements of these enzymes (6). Some bacteria also show iLDH redundancy, i.e. the presence of more than one enzyme that can act on a given isomer, which is particularly prevalent among species that produce iLDHs that act on L-lactate (“L-iLDH”s) (**Supplemental File 1**) (6). This type of redundancy can make biological sense if redundant genes differ in their spatial or temporal regulation, and are thus specialized for distinct environmental contexts (7–10).

Bacteria of the genus *Pseudomonas* are found in aquatic and terrestrial settings and are common colonizers of eukaryotic hosts (11). One outstanding feature of the pseudomonads is their ability to use a wide range of carbon sources and their preferential use of organic acids, including lactate, over sugars (12). The opportunistic pathogen *P. aeruginosa* is a common cause of hospital-acquired infections and infections in immunocompromised individuals (13). *P. aeruginosa* is able to grow using both D- and L-lactate, and our group has previously shown that it produces functionally redundant L-iLDHs (**Figure 1A**) (14). Both of these enzymes, called LldD and LldA, are predicted to bind a flavin mononucleotide (FMN) cofactor and to couple L-lactate oxidation to reduction of the quinone pool (15). The maintenance of redundant L-iLDHs in *P. aeruginosa*, which is unusual among the pseudomonads in that it can thrive in natural environments and in human infection sites (16), raises the question of whether context-dependent production of L-iLDHs contributes to its adaptability.

**Figure 1.**
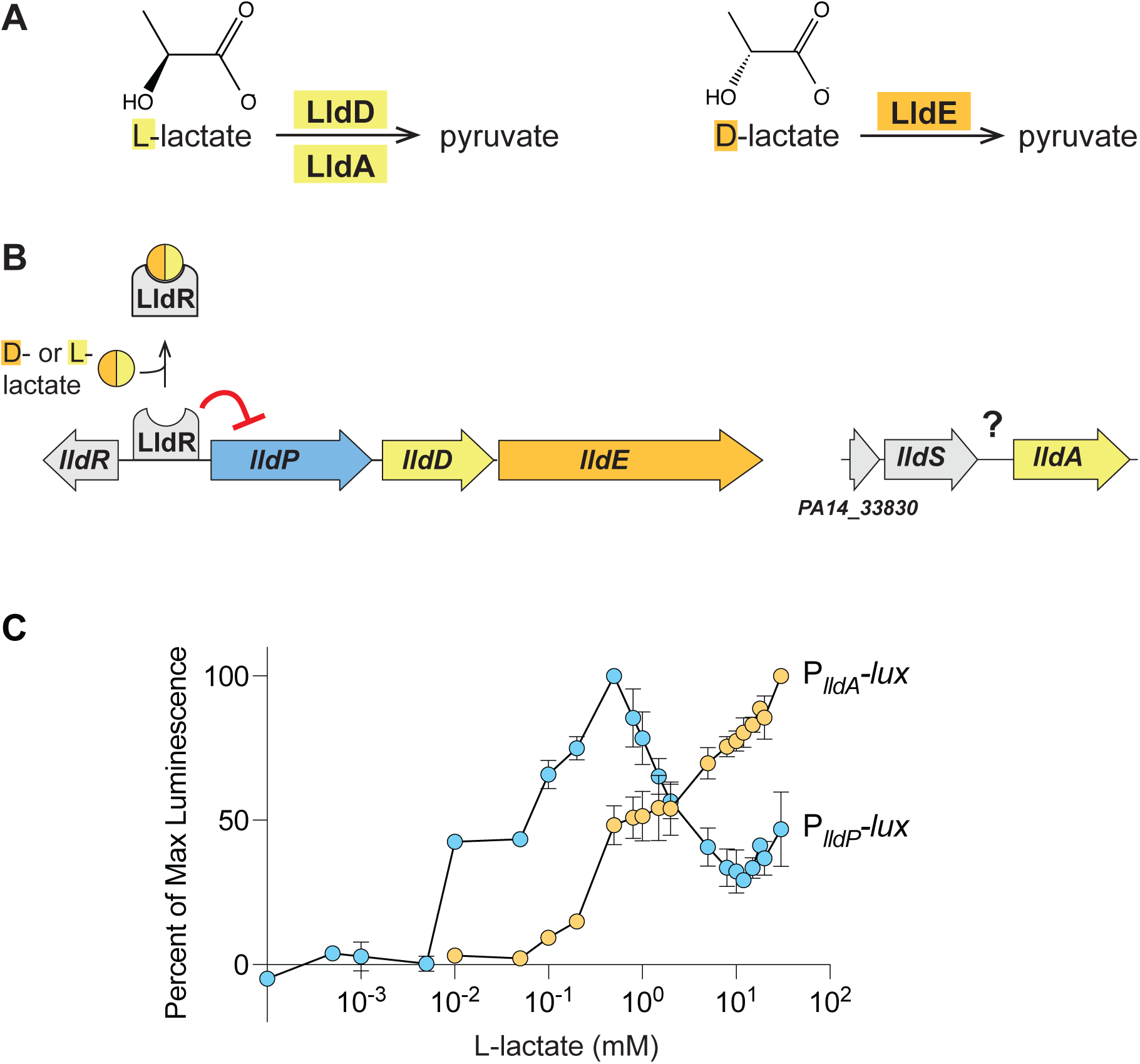
Genes for *P. aeruginosa* L-lactate dehydrogenases are sensitive to different L-lactate isoforms and concentrations. (**A**) Reactions carried out by the L-lactate dehydrogenases (LldD, LldA) and the D-lactate dehydrogenase (LldE). (**B**) Chromosomal arrangement of genes associated with lactate utilization in *P. aeruginosa*. The *lldR* gene encodes a transcriptional repressor for the *lldPDE* operon, while details regarding the regulation of *lldA* are unknown. (**C**) Activities of the *lldA* and *lldPDE* promoters at L-lactate concentrations ranging from 0.1 µM to 30 mM. Cultures of luminescent reporter strains (P*_lldA_*-*lux* and P*_lldP_*-*lux*) were grown shaking in a 96-well plate at 37°C for 24 hours in a base medium of MOPS with 20 mM succinate. Each value shown represents the maximum luminescence produced during growth in the indicated L-lactate concentration, normalized to the maximum luminescence value produced in the most-stimulatory L-lactate concentration. Values shown for each concentration are averages of two biological replicates and error bars represent standard deviation.

In addition to its metabolic versatility and ability to colonize humans, *P. aeruginosa* is notorious for its formation of biofilms: aggregates of microbial cells encased in a self-produced and protective matrix (17). These multicellular structures are well-suited to the study of condition-dependent gene expression because they contain steep chemical gradients that lead to the formation of microniches and metabolically differentiated subpopulations (18). Prior observations by our group have suggested that *P. aeruginosa lldD* is heterogeneously expressed in laboratory-grown biofilms (14, 19) while a study that applied a machine-learning approach to transcriptomic data identified *lldA* as a locus specifically associated with human infection (20). In this work, we set out to identify conditions and mechanisms that control L-lactate dehydrogenase gene expression in *P. aeruginosa.* Our findings illustrate the complexity of redundant gene utilization, which has consequences for in vitro growth and facilitates bacterial survival during infection. Moreover, our results show that differential regulation of redundant genes translates into physiological heterogeneity during multicellular *P. aeruginosa* growth. This work provides insight into context-dependent differentiation that may contribute to *P. aeruginosa*’s success as a prominent cause of biofilm-based and chronic infections.

## Results

### *lldPDE* and *lldA* are differentially expressed in response to isomer identity and L-lactate concentration

The genes for the L-lactate dehydrogenases LldD and LldA are situated at distinct sites on the *P. aeruginosa* chromosome. *lldD* is located within an operon that also codes for a lactate permease (LldP) and a D-lactate dehydrogenase (LldE, **Figure 1A**), while *lldA* is monocistronic (**Figure 1B**). Previously we showed that both lactate isomers induce *lldPDE* expression, while only L-lactate induces *lldA*, and that *lldPDE* and *lldA* exhibit different expression dynamics in liquid cultures (14). These results are consistent with the idea that differential expression might allow the redundant i-LDHs to contribute to fitness under distinct environmental conditions. Accordingly, we sought to examine the parameters and investigate the roles of potential regulators that might differentially affect *lldD* and *lldA* expression.

First we asked if expression of *lldD* and *lldA* is induced by different L-lactate concentrations. To address this question, we engineered the reporter strains P*_lldP_*-*lux* and P*_lldA_*-*lux*, which express the *luxCDABE* operon under control of the indicated promoters (21). *lux*-based reporter constructs make the luciferase enzyme (LuxAB) as well as its substrate (via LuxCDE) and have increased sensitivity to promoter activity compared to fluorescent protein-based reporters (22). In the P*_lldP_*-*lux* and P*_lldA_*-*lux* strains, the constructs are cloned into a neutral site on the chromosome; each luciferase signal serves as a readout for promoter activity and therefore reports *lldPDE* or *lldA* expression. We grew planktonic cultures of the P*_lldP_*-*lux* and P*_lldA_*-*lux* reporter strains with 20 mM succinate and various concentrations of L-lactate ranging from 0.1 µM to 30 mM. We detected luminescence from P*_lldP_*-*lux* expression already at 10 µM L-lactate, while that of P*_lldA_*-*lux* required 10x higher concentrations. In contrast to P*_lldA_*-*lux*, whose expression positively correlated with the L-lactate concentration, P*_lldP_*-*lux* expression peaked at 500 µM L-lactate (**Figure 1C**). These findings indicate that expression of *lldPDE* and *lldA* are fine-tuned to different L-lactate concentrations and suggest distinct regulatory mechanisms.

### LldS is required for *lldA* expression

Although previous studies have identified a transcriptional repressor called LldR that controls expression of *lldPDE* (23), regulators of *P. aeruginosa lldA* expression have not yet been described. To test whether LldR impacts *lldA* expression, we deleted *lldR* in a P*_lldA_*-*gfp* reporter strain (which contains a P*_lldA_*-driven GFP expression construct cloned at a neutral site on the chromosome), and grew this strain in liquid medium with L-lactate. Deletion of the *lldR* gene had only a modest effect on P*_lldA_* activity (**Figure S1**), indicating that LldR is not a major regulator of *lldA* expression and supporting our finding that *lldA* and *lldD* are differentially regulated (**Figure 1C**).

In bacterial genomes, including that of *P. aeruginosa*, it is common for the gene encoding a transcriptional regulator to lie adjacent to a target of the regulator (24). The gene just upstream of *lldA*, *PA14_33840*, which we named *lldS* (**Figure 1B**), encodes a regulatory protein predicted to contain a LysR-type substrate binding domain and a DNA-binding domain (25). LysR-type transcription factors have been implicated in the regulation of lactate utilization in other bacteria (6, 26). To test whether PA14_33840 affects *lldA* expression, we deleted *PA14_33840* in a Δ*lldD* mutant. This strain was unable to grow when L-lactate was provided as a sole carbon source, recapitulating the phenotype of a Δ*lldD* Δ*lldA* mutant (14), but did not show a growth defect when succinate was provided as the sole carbon source (**Figure 2A**). Moreover, we deleted *PA14_33840* in the P*_lldA_*-*gfp* reporter strain background and found that this abolished P*_lldA_* activity in the presence of L-lactate (**Figure 2B**). Chromosomal complementation (insertion of the wild-type *PA14_33840* allele at the native site in Δ*PA14_33840*) restored *lldA* expression to wild-type levels. Together, these observations suggest that *PA14_33840* encodes a transcriptional regulator that directly activates P*_lldA_* expression in response to L-lactate (**Figure 2C**). We have therefore named it LldS (**Figure 2C**).

**Figure 2.**
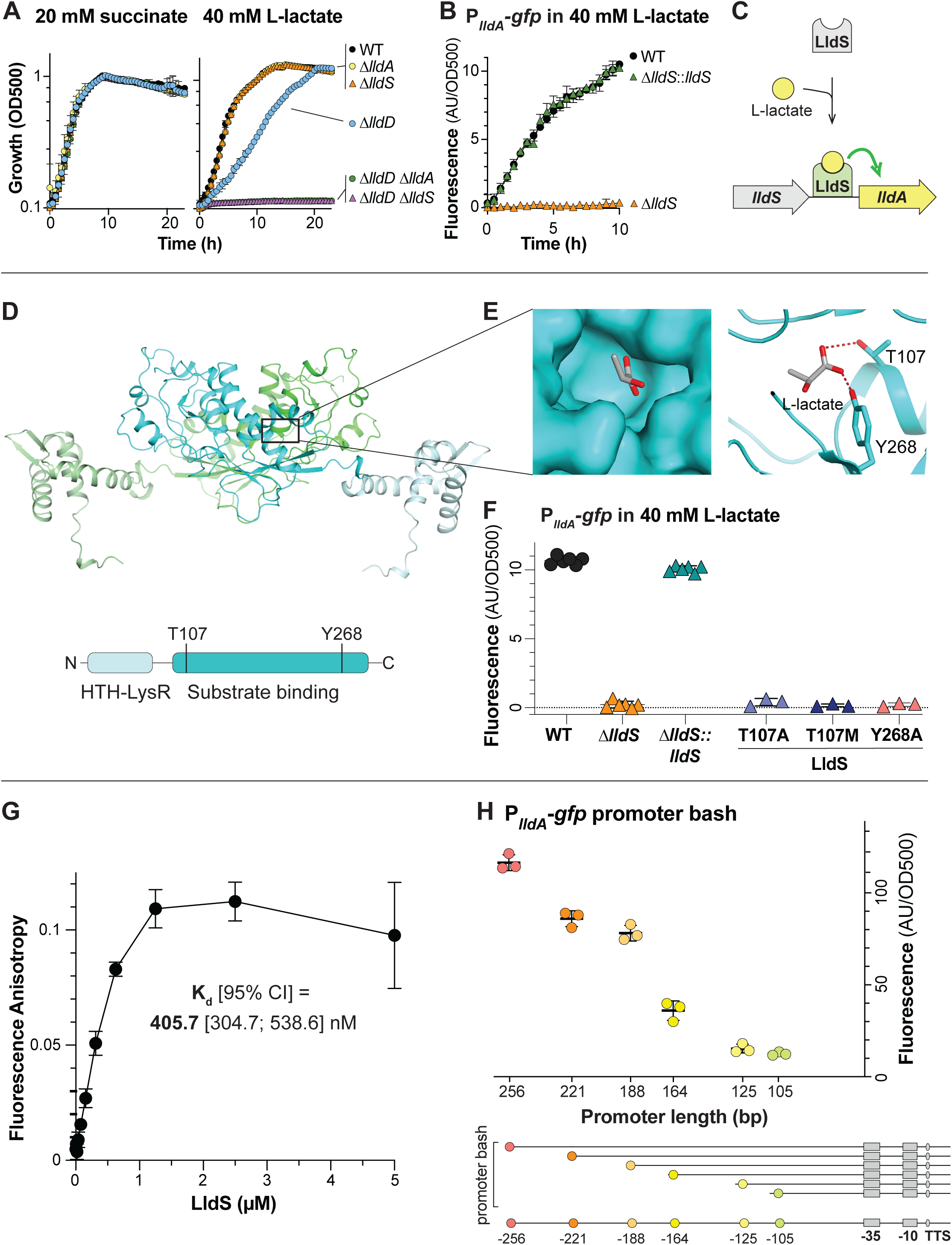
LldS (PA14_33840) is necessary for expression of *lldA* and likely senses L-lactate via its ligand-binding domain. (**A**) Growth of WT and mutants, lacking various genes associated with lactate metabolism, as liquid cultures in MOPS medium containing 20 mM succinate (left) or 40 mM L-lactate (right) as the sole carbon source. (**B**) *lldA* promoter activity in liquid cultures of WT, Δ*lldS*, and the Δ*lldS* complementation strain grown in MOPS medium containing 40 mM L-lactate. We note that growth of Δ*lldS* under this condition is supported by LldD, as indicated by the results in panel (A). (**C**) Schematic of the proposed mechanism regulating *lldA* expression. (**D**) (Top) AlphaFold-predicted structure of LldS dimer with the individual monomers colored green and cyan and with lighter shades representing the DNA-binding domains. (Bottom) Domain architecture of the LldS protein. (**E**) Left: Molecular surface of the predicted LldS binding pocket, containing L-lactate. Right: Ribbon model of the predicted LldS binding pocket with two residues, T107 and Y268, shown interacting with L-lactate. (**F**) *lldA* promoter activity in liquid cultures of Δ*lldS* strains complemented with wild-type *lldS* or with the LldS point mutants T107A, T107M, and Y268A. Strains were grown in MOPS medium containing 40 mM L-lactate. Fluorescence values were taken 5-6 hours after the onset of stationary phase. Data points represent biological replicates and error bars represent standard deviation. (**G**) Binding curve of LldS to a 5’FAM-labeled DNA probe containing 256 bp upstream of the start codon of *lldA*. Protein concentration ranged from 9.3 nM to 5 µM and the probe concentration was 5 nM. The calculated K_d_ value is shown, and error bars represent standard deviation of 2-3 replicates per concentration. (**H**) Top: Fluorescence of P*_lldA_*-*gfp* reporter strains with promoter regions of the indicated length. Each value was normalized by subtracting the average background fluorescence value of Δ*lldS* containing the full-length promoter construct. Cultures were incubated for 15 hours in MOPS medium containing 40 mM L-lactate. Data points represent biological replicates and error bars represent standard deviation. Bottom: Diagram depicting the truncations made for “promoter bash” constructs. The predicted -10 and -35 boxes and transcription start site (TSS) are indicated. These motifs were identified using the SAPPHIRE tool (89). For plots shown in panels A and B, error bars represent the standard deviation of biological triplicates and are obscured by the point marker in some cases.

To further evaluate the potential of LldS as an L-lactate-dependent transcriptional activator, we used AlphaFold2 (27) to predict its structure and identified a potential L-lactate binding pocket based on the structure of an LldS homolog in complex with a peptide ligand that has a D-alanine at the C-terminus (PDB 4WKM) (28) (**Figure 2D**). This pocket contains two residues that might coordinate the carboxyl group of L-lactate, T107 and Y268 (**Figure 2E**), which are equivalent to the interactions for the carboxyl group of D-alanine. To test whether these residues contribute to L-lactate-dependent expression of *lldA*, we generated the LldS point mutants T107A, T107M, and Y268A. We predicted that conversion of the polar threonine and tyrosine residues to alanine would interfere with hydrogen bonding to L-lactate, and that the conversion of threonine to methionine would sterically hinder L-lactate in the binding pocket. We found that all three mutations prevented activation of P*_lldA_*in response to L-lactate (**Figure 2F**).

### LldS binds the *lldA* promoter sequence

To examine the interaction between LldS and the sequence upstream of *lldA*, we purified 6x-histidine-tagged PA14 LldS protein from *E. coli* and assayed its binding to a 5’-fluorophore-labeled DNA composed of the 256-bp sequence upstream of *lldA* (i.e. the putative *lldA* promoter region) using a fluorescence polarization experiment. We generated binding curves by incubating 5 nM of the DNA with LldS at concentrations ranging from 9.37 nM to 5 µM, and determined K_d_ values for the DNA-protein interaction (**Figure 2G**). We calculated an average K_d_ [95% CI] value of 405.7 [304.7; 538.6] nM for LldS probe binding.

Having found that LldS binds to the 256 bp upstream of *lldA*, we sought to further narrow down the sequence required for *lldA* expression. We therefore characterized the region upstream of this gene using a “promoter bash” approach (29). P*_lldA_*-*gfp* reporter strains with promoters of different lengths (256, 221, 188, 164, 125, and 105 bp) were grown as liquid cultures in medium containing L-lactate. We observed a gradual decrease in P*_lldA_*-*gfp* activity with decreasing promoter length. However, we observed the most pronounced difference in P*_lldA_*-*gfp* activity between the constructs that contained 164 versus 188 bp of the sequence upstream of the start codon, which may indicate that a transcription factor binds in this region (**Figure 2H**).

### *lldPDE* and *lldA* respond differentially to ◻-hydroxycarboxylates

To test whether metabolites other than D/L-lactate affect P*_lldP_*or P*_lldA_* activity, we screened 95 compounds using plate PM1 (Biolog, Inc.) (**Supplemental File 2**). A dual-fluorescent transcriptional reporter strain, containing the *lldA* promoter driving *mScarlet* expression and the *lldPDE* promoter driving *gfp* expression, was used for these screens (**Figure S2**). In this reporter strain, the constructs are cloned into a neutral site on the chromosome; each fluorescence signal serves as a readout for promoter activity and therefore reports *lldPDE* or *lldA* expression. To screen for activating compounds, we tested for increased expression in a base medium containing succinate as the sole carbon source. For *lldPDE* expression we did not identify any activating compounds other than L-lactate (plate PM1 does not contain D-lactate), but for *lldA* expression we found that, in addition to L-lactate, alpha-hydroxybutyrate (α-HB) had a significant stimulatory effect (**Figure S2**). To screen for inhibitory compounds, we tested for decreased expression in a base medium containing succinate and L-lactate. For *lldA* expression we did not identify any inhibitory compounds, but for *lldPDE* expression we found that glycolate, which differs from lactate by the removal of one methyl group, had a strong inhibitory effect.

To verify the screen results, we surveyed the effects of α-HB, glycolate, or D-lactate on *lldA* (**Figure 3A**) and *lldPDE* expression (**Figure 3B**) by adding the compounds at concentrations ranging from 0.2 to 10 mM to a base medium containing either succinate alone or succinate with L-lactate. In cultures grown in a base medium containing succinate, we confirmed that α-HB was the only compound of the three that affected *lldA* expression and that this effect was abolished in the Δ*lldS* background. This suggested that LldS activity might be affected by both L-lactate and α-HB; we therefore examined the activity of the more-sensitive P*_lldA_*-*lux* reporter in media containing a range of concentrations of each compound and found that the promoter is more sensitive to L-lactate (**Figure S3**). *lldPDE* expression, meanwhile, was affected by all three surveyed compounds: D-lactate showed the expected stimulatory effect, α-HB showed a moderate stimulatory effect, and glycolate slightly inhibited even background expression. This background is residual expression detected in the absence of an exogenous stimulant and may arise from endogenous production of D-lactate. (*P. aeruginosa* can produce D-lactate via the enzyme LdhA (30); it does not produce L-lactate.) The glycolate effect is likely to be LldR-mediated, as its inhibitory function is lost in a Δ*lldR* strain (**Figure 3B**), which has high lactate-independent *lldPDE* expression.

**Figure 3.**
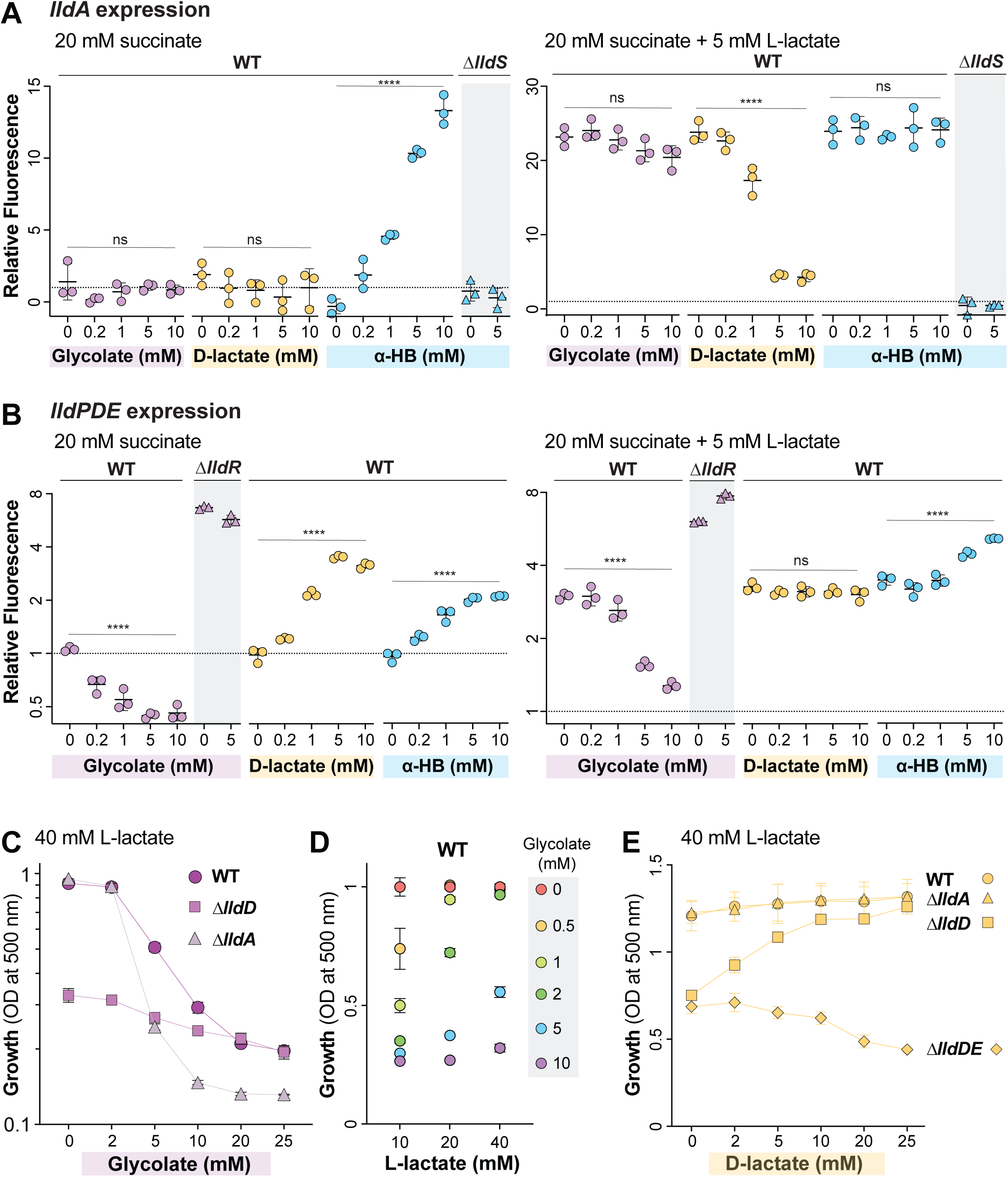
Identification of additional effectors of lactate gene regulation. (**A,B**) Fluorescence values 5-6 hours after the onset of stationary phase produced by the *lldA* and *lldPDE* reporter strains. The dashed line represents fluorescence of a wild-type culture grown with 20 mM succinate and fluorescence values are expressed in relation to this value. Cultures were grown in liquid MOPS medium containing 20 mM succinate (left graphs) or 20 mM succinate + 5 mM L-lactate (right graphs) and amended with the indicated compounds. Fluorescent values of mutant strains (Δ*lldS* for the *lldA* graphs and Δ*lldR* for the *lldPDE* graphs) are also shown for select compounds as indicated. Each dot is representative of a biological replicate and error bars represent standard deviation. **** indicates a p-value <0.0001; ns = not significant. (**C**) Growth at 10 hours of WT, Δ*lldD*, and Δ*lldA* grown in MOPS medium containing 40 mM L-lactate with glycolate concentrations ranging from 0-25 mM. (**D**) Growth of WT in MOPS medium containing L-lactate concentrations of 10, 20, or 40 mM with added glycolate ranging from 0 to 10 mM. Measurements were taken at the time point when the 0 mM glycolate condition for each L-lactate concentration reached stationary phase. Values shown are relative to growth yield at 0 mM glycolate for each respective L-lactate concentration. (**E**) Growth at 15 hours of WT, Δ*lldD*, Δ*lldA*, and Δ*lldDE* strains grown in MOPS medium containing 40 mM L-lactate with D-lactate concentrations ranging from 0-25 mM.

In the “stimulatory” base medium containing both succinate (20 mM) and L-lactate (5 mM), the addition of glycolate had no significant effect on *lldA* expression. In contrast, increasing amounts of D-lactate had an inhibitory effect on *lldA* expression, beginning at 1 mM. We also observed that, in the stimulatory base medium, α-HB had no effect on *lldA* expression. This is consistent with our results suggesting that P*_lldA_* is more sensitive to L-lactate than α-HB (**Figure S3**) and indicates that α-HB is not a strong competitor for LldS binding when L-lactate is present. With respect to *lldPDE* expression, we found that glycolate showed an inhibitory effect even when L-lactate was present, particularly when it was added at an equimolar concentration. D-lactate did not stimulate *lldPDE* expression beyond the level arising from L-lactate-dependent induction. Finally, α-HB showed a slight stimulatory effect comparable to that observed in medium without lactate.

Overall, the results of these experiments show that the activities of P*_lldP_* and P*_lldA_* are differentially affected by various alpha-hydroxy acid compounds. While *lldPDE* expression is stimulated by D-lactate and inhibited by glycolate, *lldA* expression is inhibited by D-lactate and relatively unaffected by glycolate. The effect of glycolate on P*_lldP_* is mediated by the repressor LldR (**Figure 3B**). The effect of D-lactate on P*_lldA_* could indicate that LldS, which is activated by L-lactate, is deactivated by the D-isomer of lactate. An alternative explanation, however, is that D-lactate appears to inhibit L-lactate-dependent *lldA* expression because less L-lactate enters the cell when both isomers are provided simultaneously, due to competition for the lactate permease (31). Finally, both promoters are activated by α-HB, but in the case of P*_lldA_* this requires the regulator LldS (**Figure 3A**), while for P*_lldP_* we do not yet have an indication as to how this activation is mediated.

Consistent with the observations that LldD is the primary contributor to growth on L-lactate under standard conditions (**Figure 2A** and (14)) and that *lldPDE* expression is inhibited by glycolate (**Figure 3B**), previous studies have shown that *P. aeruginosa* growth on lactate is inhibited by the addition of glycolate (23, 32). This raised the question of whether growth of *P. aeruginosa* on L-lactate plus glycolate increases its reliance on LldA, whose expression is unaffected by glycolate (**Figure 3A**). To test this, we grew PA14 WT alongside Δ*lldD* and Δ*lldA* mutants in 40 mM L-lactate with added glycolate ranging from 0 to 25 mM (**Figure 3C**). As expected, we found that glycolate inhibited wild-type growth, causing a sharp decrease in growth yield at 5 mM glycolate. In contrast, although Δ*lldD* showed decreased growth consistent with LldD’s primary role in L-lactate utilization, the effect of increasing glycolate on this yield was less pronounced and was indistinguishable from WT at concentrations of 20 mM and higher. The residual growth of WT and Δ*lldD* at high glycolate concentrations indicates LldD-independent L-lactate utilization that is less sensitive to glycolate. Accordingly, in contrast to Δ*lldD*, the Δ*lldA* mutant showed a pronounced decrease in growth yield at glycolate concentrations of 5 mM and higher when compared to the WT. These results show that LldA makes an important contribution to growth of *P. aeruginosa* on L-lactate when glycolate is present. Finally, to test whether sensitivity to glycolate is affected by the concentration of L-lactate provided, we performed similar experiments with a base L-lactate concentration of 10 or 20 mM for a direct comparison with the effects of 40 mM L-lactate on growth (**Figure 3D**). We found that cultures grown on lower concentrations of L-lactate were sensitive to lower concentrations of glycolate, with glycolate negatively affecting growth starting at only 0.5 mM glycolate in the 10 mM-L-lactate culture and at 2 mM glycolate in the 20 mM-L-lactate culture. This observation suggests that glycolate might competitively bind to LldR and enhance its repression of *lldPDE* expression.

In addition to the negative effect of glycolate on P*_lldP_*activity, our expression analysis revealed an inhibition of P*_lldA_* activity by D-lactate (**Figure 3A)**. We hypothesized that this would lead to an increased reliance on LldD under conditions where both lactate isomers were present. We therefore predicted that addition of D-lactate would be detrimental to the growth of the Δ*lldD* mutant on L-lactate. We grew WT *P. aeruginosa* alongside Δ*lldD* and Δ*lldA* mutants in 40 mM L-lactate with added D-lactate concentrations ranging from 0 to 25 mM. We found that added D-lactate had no effect on WT and Δ*lldA* growth. Unexpectedly, however, it had a stimulatory effect on Δ*lldD* growth (**Figure 3E**). We reasoned that this stimulation arose from the utilization of D-lactate via LldE activity (**Figure 1A**), and therefore repeated the experiment, this time including a Δ*lldDE* mutant. As expected, this mutant showed decreased growth in D-lactate concentrations of 10 mM or higher.

### The genomic context of *lldA* suggests a link to iron availability

Having observed effects of organic metabolites and local regulators on expression of *P. aeruginosa*’s redundant L-iLDH genes, we next examined the phylogenetic relationship of these proteins for clues regarding their respective physiological roles. Instead of or in addition to LldD and/or LldA, some organisms possess an L-iLDH encoded by a three-gene cluster referred to as *lutABC* (33). We searched for *lldD*, *lldA* and *lutABC* homologs in *Pseudomonas* genomes from the Pseudomonas Genome DB (34), choosing one representative strain genome for each *Pseudomonas* species. Out of the 213 strains with L-iLDHs, we identified LldD and LldA homologs in 179 of these genomes. **Figure 4A** shows a phylogenetic tree generated using the corresponding sequences and depiction of their gene arrangements. **Supplemental File 1** lists all 213 analyzed strains and indicates their L-iLDH profiles; those containing LldA or LldD are arranged according to the Lld phylogeny. **Figure 4B** shows the total numbers of analyzed genomes that contain each of the indicated L-iLDH gene arrangements and the numbers of genomes that contain two or more L-iLDH homologs. In the tree shown in **Figure 4A**, the *lldD* and *lldA* homologs formed independent clusters. Each of these clusters, however, also contained subgroupings that largely correlated with the genomic neighborhood/arrangement of *lldD* and *lldA* genes. *lldD* genes separated into two main subgroups: (i) those contained within an *lldPDE* operon, adjacent to a divergently transcribed *lldR* homolog (indicated in blue; as in *P. aeruginosa* PA14); and (ii) those that were monocistronic, located next to a divergently transcribed gene for a LysR-family transcriptional regulator (indicated in orange) (**Figure 4A**).

**Figure 4.**
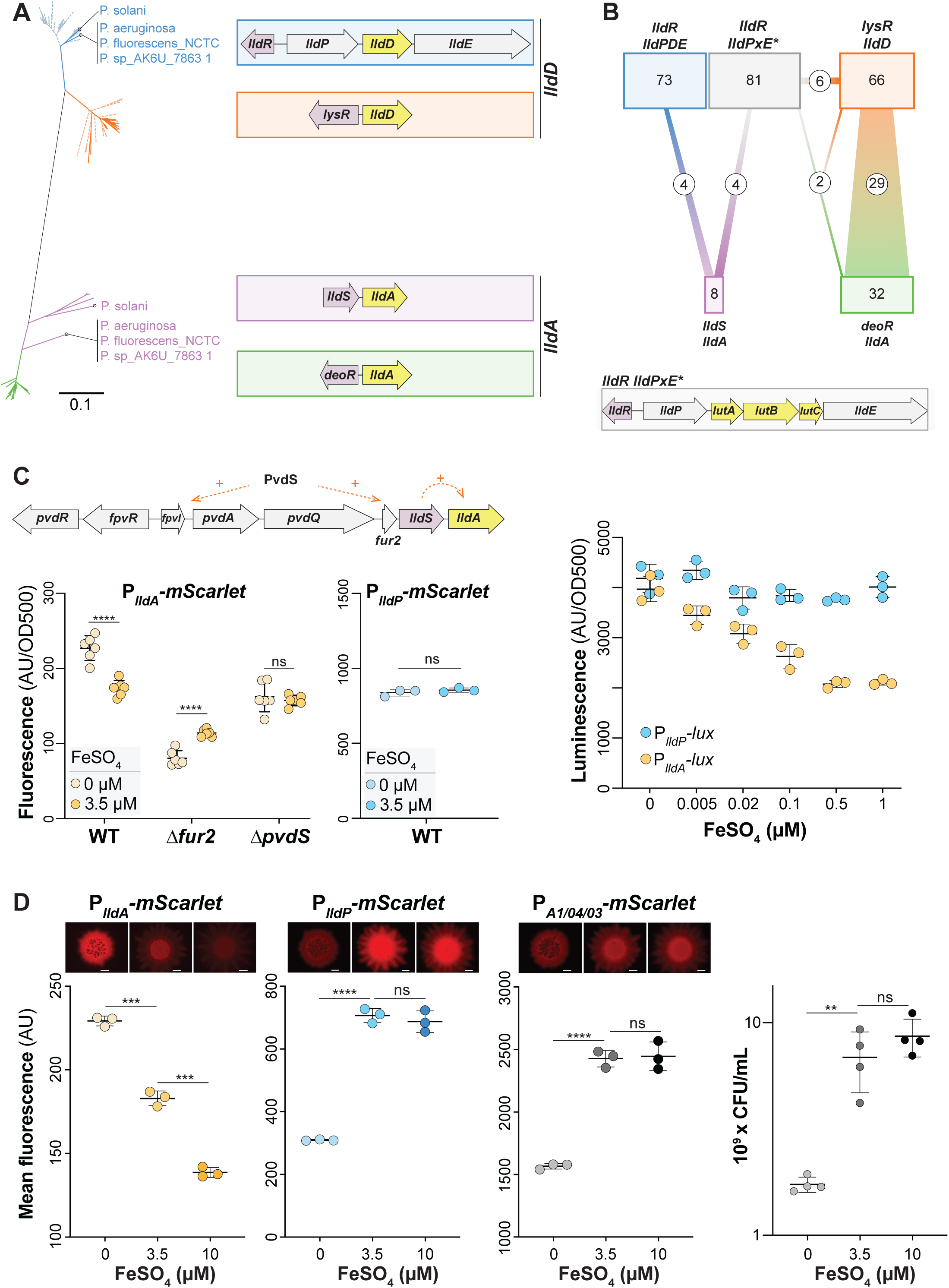
*lldA*, but not *lldPDE*, expression is sensitive to iron availability. (**A**) Phylogenetic tree of *lldD* and *lldA* genes obtained from 213 pseudomonad genomes. Only one representative strain per species was included, unless strains showed different *lld* arrangements. Blue and orange lines represent genes with homology to *lldD* while green and purple lines represent genes with homology to *lldA*. Dotted lines indicate strains with one L-iLDH; solid lines indicate strains with more than one. The color of each cluster corresponds to the outlines surrounding the gene arrangement patterns in the panel on the right. Species/strain name is written if the strain has two L-iLDH genes. Only eight *Pseudomonas* species (including *P. aeruginosa*) possess the *lldS-lldA* gene arrangement outlined in purple. (**B**) Diagram depicting genome associations for the arrangement patterns shown (A) (indicated by a consistent color coding). An additional gene locus with the L-lactate utilization genes *lutABC*, which is not reflected in the tree, is outlined in gray and depicted at the bottom of this panel. The number of species possessing each gene arrangement is indicated in its corresponding box, and the connections represent the number of species with multiple L-iLDHs and their corresponding gene arrangements. (**C**) Top: In *P. aeruginosa*, *lldS-lldA* lies directly downstream of several genes involved in iron regulation and uptake. Left: *lldA* promoter activity in liquid cultures of WT, Δ*fur2*, and Δ*pvdS* strains grown in MOPS medium containing 40 mM L-lactate with ferrous sulfate added (3.5 µM) or no iron added (0 µM). Fluorescence values were taken 5-6 hours after the onset of stationary phase. Center: *lldPDE* promoter activity under the same conditions used for the P*_lldA_* reporter. Right: *lldA* and *lldPDE* promoter activity assayed using luciferase reporters. Cultures were grown in MOPS liquid medium with 40 mM L-lactate and ferrous sulfate was added as indicated. The maximum luminescence value for each condition is shown for each reporter. (**D**) Representative top-down fluorescent biofilm images and quantification of the average fluorescence across the width of the center of the biofilm for *lldA* reporter (left), *lldPDE* reporter (center left), and constitutive *mScarlet* (center right) strains. Biofilms were grown on MOPS medium with 20 mM succinate and 10 mM L-lactate, amended with ferrous sulfate as indicated. Scale bars = 2 mm. Right: Quantification of colony forming units of biofilms grown under each iron-amendment condition. Each dot is representative of a biological replicate and error bars represent standard deviation. **p<0.01, ***p<0.001, ****p<0.0001, ns = not significant.

*lldA* homologs showed a more scattered phylogeny overall, but also formed two subgroups with consistent genomic arrangements: (i) a monocistronic *lldA* gene located next to a divergently transcribed gene for DeoR-family transcriptional regulator (indicated in green) and (ii) a much less common *lldA* gene located directly downstream of an *lldS* homolog (indicated in purple) (**Figure 4A,B**). The *P. aeruginosa lldA* sequence fell within the latter subgroup. *P. aeruginosa*, however, stood out as the only species in which *lldS* and *lldA* are situated near a large chromosomal region involved in the uptake of iron (**Figure 4C**). *PA14_33830* (“*fur2*”), which lies upstream of *lldS*, bears homology to the global iron regulator Fur, is highly upregulated under low iron conditions (35, 36), and has been implicated in the regulation of a wide range of genes including those involved in iron uptake and siderophore production (37).

### Expression of *lldA* is enhanced under low-iron conditions

Given the proximity of *lldS* and *lldA* to the cluster of iron-related genes, we tested the effect of iron availability on *lldA* expression. The defined medium that we typically use for *P. aeruginosa* growth contains iron added as freshly prepared ferrous sulfate at a concentration of 3.5 µM. We grew the P*_lldA_*-*mScarlet* strain in this liquid medium with L-lactate as the carbon source, with or without the added iron. Because iron chelators were not added to the “low-iron” medium, residual amounts of contaminating iron enabled *P. aeruginosa* to grow (albeit at levels that were significantly lower than that observed for the standard (i.e. iron-amended or -replete) medium) (**Figure S4**). We found that OD-corrected *lldA* expression was enhanced under the low-iron condition when compared to expression in the standard medium (**Figure 4C, left**). This effect was not observed for *lldPDE* expression (**Figure 4C, center**). Increasing the concentration of added iron to 10 µM also had no effect on *lldA* expression under these (liquid-culture) conditions (**Figure S5**). To better visualize subtle effects on gene expression that might arise under various conditions of iron availability, we once again utilized the luciferase reporters (P*_lldA_*-*lux* and P*_lldP_*-*lux*) and tested concentrations of ferrous iron sulfate ranging from 5 nM to 1 µM. While *lldA* expression decreased in a stepwise fashion as we increased iron concentrations, we did not observe significant changes in *lldPDE* expression (**Figure 4C, right**).

How is the effect of iron on *lldA* expression mediated? To begin to address this question, we tested the contributions of two regulatory genes, *fur2* and *pvdS*, to *lldA* expression under conditions of added or omitted ferrous iron sulfate. PvdS is a sigma factor that directly controls expression of several loci in the region adjacent to *lldS*, including *fpvI* (**Figure 4C**), and that has been implicated in regulation of *lldS* (*35, 38*). We found that, in both the Δ*fur2* and Δ*pvdS* strain backgrounds, the induction of *lldA* expression observed in the WT under low-iron conditions was abolished (**Figure 4C**). These results suggest that both Fur2 and PvdS contribute to the iron-dependent effects on *lldA* expression.

Nutrient limitation is a key determinant of physiological status in biofilms because multicellularity promotes chemical gradient formation (18). We therefore tested the effect of iron availability on *lldA* and *lldPDE* expression in biofilms. Our group studies the physiology of bacteria in biofilms using a macrocolony assay, in which a suspension of cells is pipetted onto an agar-solidified growth medium and incubated in a standard atmosphere at high humidity for several days (39, 40).

To examine the effects of iron availability on *lld* expression in biofilms, we grew the P*_lldA_*- and P*_lldP_-mScarlet* reporter strains in the macrocolony assay on a defined medium containing L-lactate with different amounts of added ferrous iron sulfate. We also included a strain containing the P*_A1/04/03_*-*mScarlet* construct; this strain constitutively synthesizes fluorescent protein and can therefore serve as a control for the effects of iron availability on overall biomass production.

Additionally, to control for effects of iron availability on growth, we homogenized biofilms and plated for CFUs. As expected, biofilms grown on medium without added iron yielded a CFU count that was approximately one order of magnitude lower than biofilms grown on medium with 3.5 or 10 µM added iron, and fluorescence levels of the P*_A1/04/03_*-*mScarlet* biofilms (imaged top-down on a fluorescent microscope) correlated with these relative CFU counts (**Figure 4D**). We observed similar changes in fluorescence for P*_lldP_-mScarlet* biofilms. Notably, in comparison to P*_A1/04/03_*-*mScarlet* and P*_lldP_-mScarlet* biofilms, P*_lldA_*-*mScarlet* biofilms showed an inverse trend. In spite of the decreased biofilm growth yield observed on medium without added iron, the P*_lldA_*-*mScarlet* reporter biofilm showed a significant increase in fluorescence on this medium when compared to the 3.5-µM-added-iron condition. Additionally, although growth yields were similar with 3.5 µM- and 10 µM-added-iron, *lldA* expression was significantly lower in the high-iron condition. In summary, our experiments examining *lld* gene expression in liquid cultures and biofilms indicate that iron limitation enhances expression of *lldA*, but not *lldPDE*, consistent with the *lldA*’s chromosomal location (near iron-related genes). In biofilm experiments specifically, we also observed that the addition of excess iron was inhibitory to *lldA* expression.

### *lldPDE* and *lldA* are differentially expressed across biofilm depth

We predicted that the opposing gradients of oxygen (from the biofilm-air interface) and other resources (from the biofilm-medium interface) in macrocolony biofilms would differentially affect the expression of *lldDPE* and *lldA*. To examine this, we grew biofilms of the dual transcriptional reporter strain (P*_lldP_*-gfp P*_lldA_*-mScarlet) for three days before preparing thin sections (**Figure 5A**) (41). For this experiment, the medium contained 20 mM succinate as the primary carbon source so that growth would not depend solely on LldD/A activity. Imaging via fluorescence microscopy revealed intriguing and robust patterns of *lldA* and *lldPDE* expression (**Figure 5B**). Most notably, the activities of P*_lldA_*and P*_lldP_* showed an inverse relationship, indicating maximal expression of *lldA* and relatively low expression of *lldPDE* at the biofilm-air interface, and maximal expression expression of *lldPDE* in a region where *lldA* expression shows a pronounced decline (30-40 µm from the biofilm-air interface). We observed a similar pattern for biofilms inoculated with an equal mixture of individual P*_lldP_*-gfp and P*_lldA_*-mScarlet or of individual P*_lldP_*-mScarlet and P*_lldA_*-gfp reporter strains (**Figure S6**). The exclusionary patterning led us to hypothesize that LldD might negatively affect expression of *lldA* and/or vice versa. To test this, we created mutants lacking the L-lactate dehydrogenase genes (Δ*lldA* and Δ*lldD*) in the background of the dual (P*_lldP_*-gfp P*_lldA_-*mScarlet) transcriptional reporter strain and examined promoter activity in biofilm thin-sections. We found that although overall *lldA* expression was enhanced in the Δ*lldD* mutant (**Figure 5C**), the characteristic expression patterns of *lldPDE* and *lldA* were unaffected by the respective gene deletions (**Figure 5B**), indicating that this tight regulatory patterning is determined by other factors.

**Figure 5.**
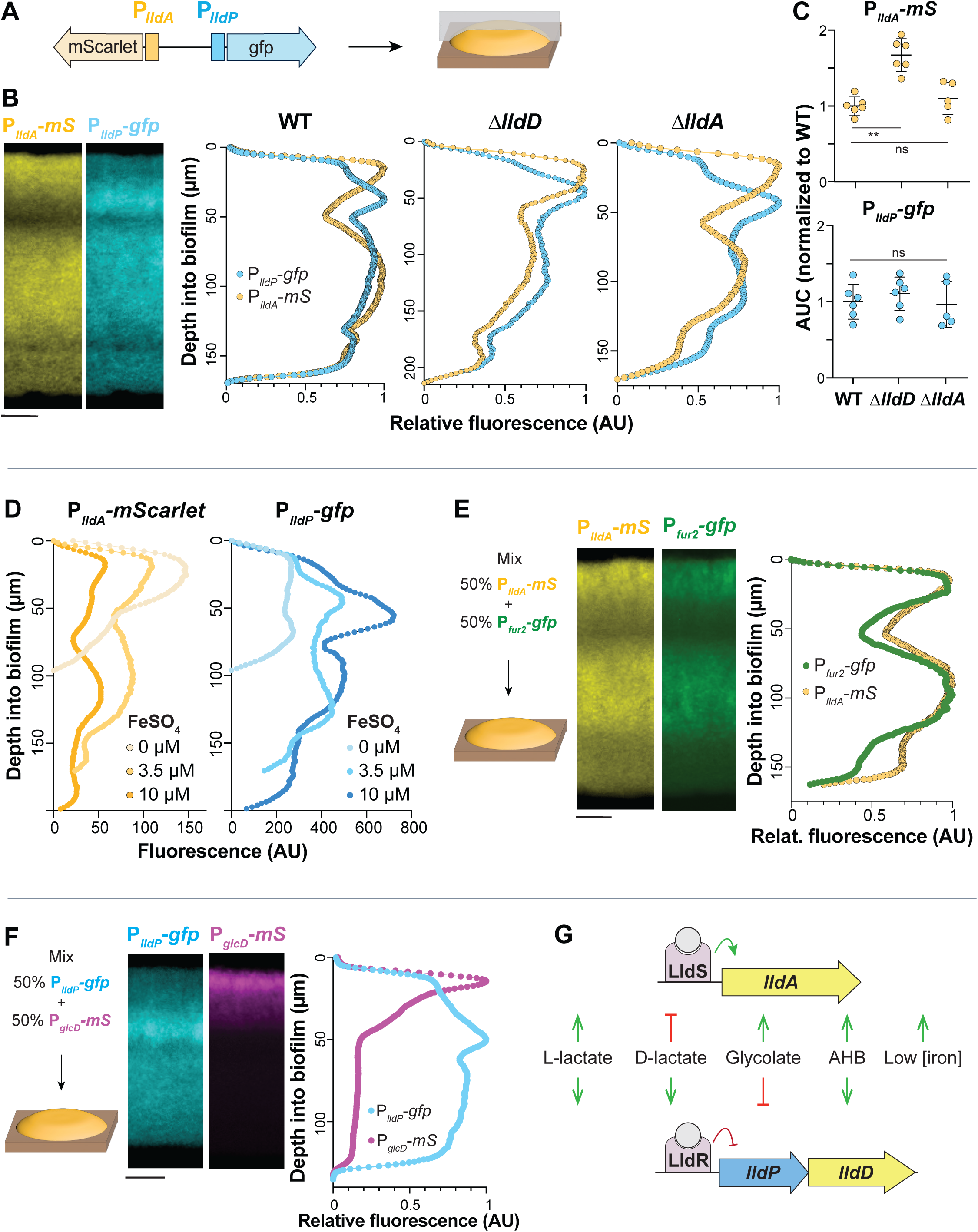
Biofilms display patterns of *lldA* and *lldPDE* expression across depth that might arise from local differences in iron availability and glycolate production. **(A)** Schematic showing the genomic arrangement of the dual reporter construct (P*_lldA_*-*mScarle*t, P*_lldP_*-*gfp)* used in these experiments and the orientation of thin-sectioning for a macrocolony biofilm. (**B**) Left: Fluorescence images of thin-section from a biofilm formed by the dual P*_lldA_*-*mScarlet* (“*mS*”), P*_lldP_*-*gfp* reporter strain. *mScarlet* fluorescence is shown in yellow and *gfp* fluorescence is shown in cyan. Right: Fluorescence across depth for the P*_lldA_*and P*_lldP_* reporters in the indicated strain backgrounds. Images and quantification are representative of at least three biological replicates. (**C**) Quantification of total dual-reporter thin-section fluorescence expressed as area under the curve (AUC) for *lldPDE* (top) and *lldA* (bottom) expression in WT, Δ*lldD*, and Δ*lldA*, normalized to average wild-type fluorescence. Each dot is representative of a single biological replicate and error bars represent standard deviation. (**D**) Fluorescence across depth for each reporter in biofilms of the dual-reporter strain grown on medium amended with ferrous sulfate as indicated. Profiles are representative of three biological replicates for each iron-availability condition. (**E**) Left: Schematic of experimental design for growth of biofilms containing two reporter strains: P*_lldA_*-*mScarlet* and P*_fur2_*-*gfp.* Center: Fluorescence images of thin-section from a mixed biofilm. *mScarlet* fluorescence is shown in yellow and *gfp* fluorescence is shown in green. Right: Fluorescence across depth for the P*_lldA_*-*mScarlet* and P*_fur2_*-*gfp* reporters. Images and quantification are representative of four biological replicates. (**F**) Left: Schematic of experimental design for growth of biofilms containing two reporter strains: P*_glcD_*-*mScarlet* and P*_lldP_*-*gfp.* Center: Fluorescence images of a thin-section from a mixed biofilm. *mScarlet* fluorescence is shown in magenta and *gfp* fluorescence is shown in cyan. Right: Fluorescence across depth for the P*_glcD_*- *mScarlet* and P*_lldP_*-*gfp* reporters. Images and quantification are representative of four biological replicates. (**G**) Visual summary of the cues that activate or inhibit expression of *P. aeruginosa lldA* and *lldPDE*. Biofilms in B, D, E, and F were grown on MOPS medium containing 20 mM succinate and 10 mM L-lactate. Scale bars = 25 µm.

The fact that resource gradients form within biofilms raised the question of whether iron limitation contributes to induction of *lldA* at the biofilm-air interface. We, therefore, sought to test whether iron availability influences the spatial patterning of *lldA* and *lldPDE* expression across biofilm depth. We grew the dual transcriptional reporter strain for three days on agar containing succinate and L-lactate and various concentrations of added iron sulfate and prepared biofilm thin sections. While we found, once again, that the general patterning of *lldA* and *lldPDE* expression was retained, we also noted that *lldA* expression increased, with decreasing iron addition, specifically in the region close to the air interface. Further, overall levels of expression varied. Iron availability affected the expression levels of both *lldA* and *lldPDE*, but in opposite ways (**Figure 5D**). With increasing concentrations of added iron (0 µM, 3.5 µM, and 10 µM), *lldA* expression levels decreased, while those for *lldPDE* increased (the latter may be due to general increase in biomass as indicated in **Figure 4D**). To further investigate iron availability across biofilm depth, we created a strain that reports expression of *fur2* (“P*_fur2_-gfp*”), which is induced by low iron conditions (35). We then grew biofilms, started from an equal mixture of P*_fur2_-gfp* and P*_lldA_*-*mScarlet*, on medium containing succinate and L-lactate and found that *lldA* and *fur2* expression were aligned across biofilm depth (**Figure 5E**). Together, these results suggest that iron availability is a significant parameter determining the pattern of *lldA* expression in biofilms.

While iron availability could explain some of the *lldA* expression features in a biofilm, it does not seem to be responsible for the distinct decrease in *lldPDE* expression close to the air interface (**Figure 5D**). We also excluded that the high *lldA* expression in this region affects *lldPDE* expression (**Figure 5B)**, indicating that its spatial patterning is influenced by other cues. Since we had found in our compound screen that glycolate inhibits *lldPDE* expression (**Figure 3A,C**), we followed up on this lead. *P. aeruginosa* contains the *glcC* gene and the adjacent *glcDEFG* operon, which are homologous to *E. coli* genes for a glycolate-sensing transcription factor that induces *glcDEFG* expression, and a glycolate oxidase complex, respectively (42). We constructed a P*_glcD_*-*mScarlet* reporter strain and confirmed using liquid-culture experiments that its activity is induced by glycolate (**Figure S7**). To test the hypothesis that glycolate contributes to the pattern of *lldPDE* expression in biofilms, we used equal mixtures of P*_glcD_*-*mScarlet* and P*_lldP_*-*gfp* to inoculate agar plates containing succinate and L-lactate (**Figure 5F**). We found that *glcD* expression was highly induced specifically at the biofilm-air interface. This finding suggests a build-up of glycolate at the top of the biofilm, which could explain the decreased *lldPDE* expression in this zone.

Having shown that iron availability and exposure to specific alpha-hydroxy carboxylates influence the differential regulation of *lldD* and *lldA* (**Figure 5G**), we next tested the contributions of their respective enzymes during macrophage infection.

### *lldD* and *lldA* contribute to persistence of *P. aeruginosa* in macrophages

Exposure of macrophages to bacteria or bacterial products has been reported to induce a “Warburg-like effect” in which macrophages exhibit increased glycolysis and decreased oxidative phosphorylation (43). Accordingly, studies have suggested that levels of lactate, a byproduct of glycolysis (44), are increased in infected macrophages and that lactate promotes growth of intracellular bacterial pathogens within these cells (45, 46). We therefore used RAW264.7 cells to test whether L-lactate dehydrogenase genes are expressed in the presence of macrophages, and whether they contribute to *P. aeruginosa* success during infection. Macrophages incubated with the PA14 P*_lldA_*-*mScarlet* reporter strain showed red fluorescence that was absent in reporterless and uninfected controls, indicating that *lldA* is expressed during infection (**Figure 6A**). To evaluate whether the L-lactate dehydrogenases contribute to *P. aeruginosa*’s success during macrophage infection, we used a gentamicin protection assay (47). We found that macrophages infected with deletion mutants lacking the L-lactate dehydrogenase genes *lldD* or *lldA*, or the mutant lacking its regulator *lldS* showed a comparable decrease in bacterial burden when compared to those infected with WT *P. aeruginosa* (**Figure 6B**). The observation that the double mutant Δ*lldD*Δ*lldA* phenotype is similar to the individual L-iLDH mutants suggests that the contributions of LldD and LldA are not additive under this condition. Nevertheless, these results indicate that both LldD and LldA contribute to *P. aeruginosa* persistence within macrophages.

**Figure 6.**
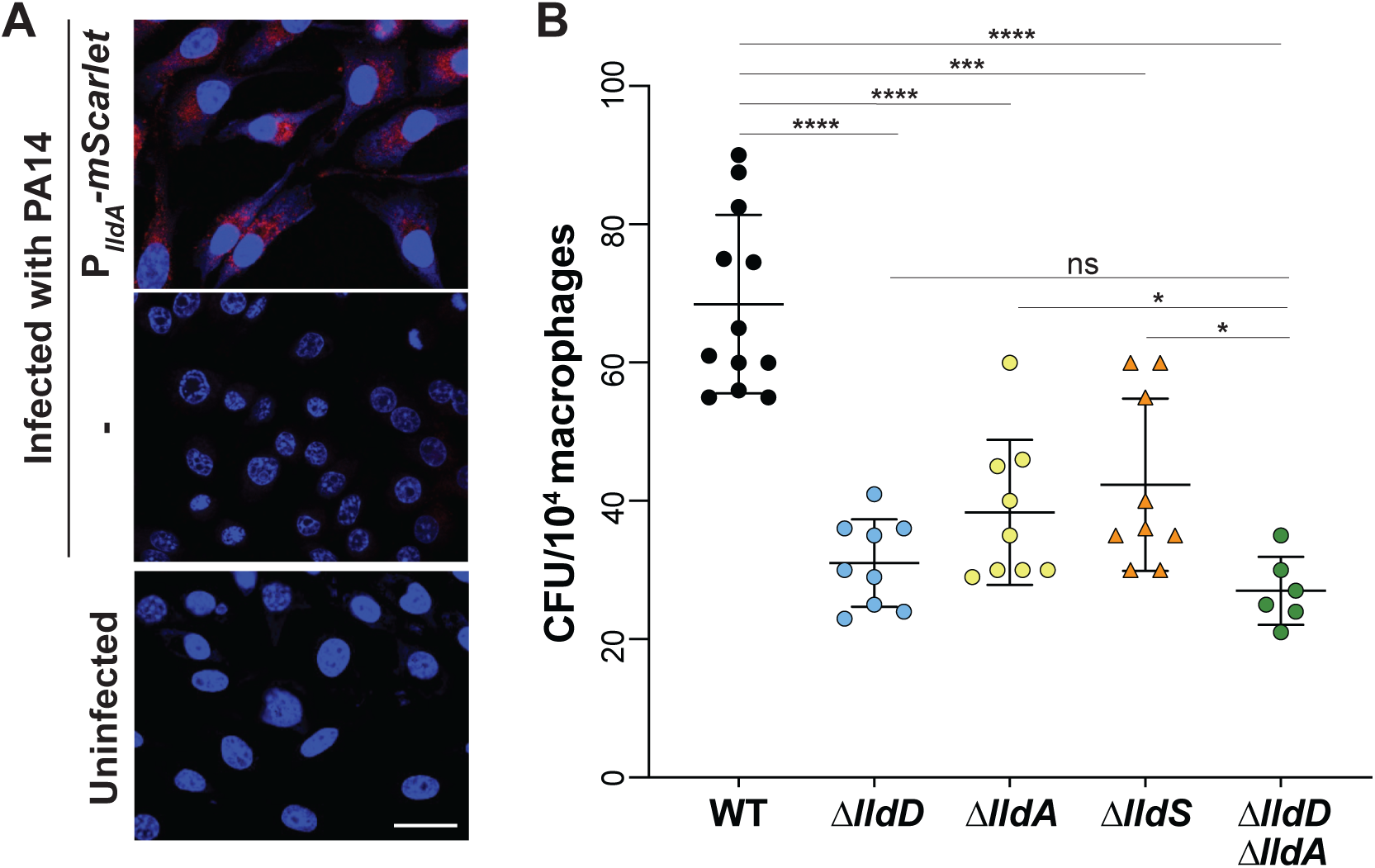
Expression of *lldA* and contributions of *lld* genes during macrophage infection. (**A**) Fluorescence images of RAW264.7 macrophages infected with WT PA14 (middle), the P*_lldA_*- *mScarlet* strain (top), and of an uninfected control (bottom). DAPI fluorescence is shown in blue and mScarlet fluorescence is shown in red. Scale bar is 10 µm. (**B**) Intracellular burden of *P. aeruginosa* WT and indicated mutant strains in RAW264.7 macrophages 3 hours post-infection and subjected to the gentamicin protection assay. Each dot represents one replicate and error bars represent standard deviation. *p<0.05, ***p<0.001, ****p<0.0001, ns = not significant.

## Discussion

Enzymes that convert L-lactate to pyruvate allow bacteria to directly link a common carbon source to central metabolism. They contribute to growth alongside fermentative bacteria and eukaryotes, and during colonization of eukaryotes. Within the human body, for example, L-lactate has been shown to accumulate to 1.5-3 mM in blood and tissue under normal physiological conditions, and up to 40 mM under inflammatory or cancerous conditions (44), making it a significant carbon source for pathogens. Studies in various bacteria, including *Neisseria gonorrhoeae*, *Staphylococcus aureus*, and *Mycobacterium tuberculosis*, support the idea that L-lactate utilization contributes to success in the host and/or to virulence (45, 48–52). Coincidentally, all of these organisms have the capacity to produce multiple L-lactate dehydrogenases.

*P. aeruginosa* is a biofilm-forming, opportunistic pathogen that harbors traits similar to those of the nonpathogenic members of the *Pseudomonas* genus in addition to adaptations that allow it to cause disease in diverse hosts, including humans. These include the ability to grow at high temperatures and the production of a broad array of virulence factors (53–55). We were intrigued by *P. aeruginosa*’s possession of a redundant L-lactate dehydrogenase gene and the fact that this gene lies adjacent to the “pvd region” of the chromosome (35, 56), which contains multiple genes involved in iron acquisition and the response to iron limitation (**Figure 4C**). This close proximity of the *lldS*-*lldA* locus and iron-related genes is unique to *P. aeruginosa* among the pseudomonad genomes that we surveyed. In our prior work, we showed that both of *P. aerugiosa*’s genes for L-iLDH enzymes–*lldD* and *lldA*–are expressed in liquid cultures in media containing L-lactate, including artificial sputum media, which contain this carbon source at millimolar concentrations. In liquid batch cultures, *lldD* is expressed before *lldA* and is sufficient to support wild-type growth dynamics (14). However, transcriptomic studies have shown that *lldA* is specifically induced in infection models and under infection conditions (20, 57, 58), which indicates an important role for LldA during association with hosts. We therefore investigated the regulation of *P. aeruginosa*’s *lldD* and *lldA* genes, the pattern of their expression in biofilms, and their contributions to host cell infection.

In this study, we confirmed that *lldD* expression is induced by both isomers of lactate (**Figure 3B**), which alleviate repression by LldR (23). We discovered that *lldA* induction is controlled by the activator LldS (**Figure 2**), which is encoded by an adjacent gene and responds specifically to the L-isomer. The chromosomal location of *lldA* led us to hypothesize that its expression could be affected by iron availability, and we indeed observed an inverse relationship between gene expression and iron availability that was unique to this L-lactate dehydrogenase gene (**Figure 4C,D**). We also found that the expression of *lldA* correlates with expression of *fur2* (**Figure 5E**), a putative regulatory gene located next to *lldS* that has previously been shown to respond to iron limitation (35, 36). Moreover, we observed reduced *lldA* expression for mutants lacking *fur2* and *pvdS* (**Figure 4B**), which codes for a sigma factor that controls a large regulon of genes induced by iron limitation, including *fur2* (35). We hypothesize that in response to low iron, PvdS increases *fur2* transcription, and Fur2 acts to increase *lldA* transcription in the presence of L-lactate, either by directly binding the *lldA* promoter or through increasing expression of *lldS*. Iron limitation may act as a proxy signal for conditions in infection sites, where host metabolites act to sequester iron and decrease its availability for pathogens (59, 60), and where L-lactate may be available as a carbon source. Accordingly, in *N. gonorrhoeae*, it has also been suggested that the induction of a L-lactate dehydrogenase gene by iron limitation is an adaptation that promotes this pathogen’s utilization of L-lactate during infection (51).

Another notable aspect of the iron-responsive regulation of *lldA* is that *P. aeruginosa*’s other L-lactate dehydrogenase gene, *lldD*, is cotranscribed with *lldE*. LldE codes for a multidomain D-lactate dehydrogenase that is predicted to contain Fe-S clusters (61). By inducing *lldA*, as opposed to *lldPDE*, under low-iron conditions, *P. aeruginosa* poises itself to utilize L-lactate but avoids production of a costly, iron-containing, and potentially superfluous enzyme.

In addition to the effect of iron availability, we evaluated the effects of a suite of metabolites on P*_lldP_* and P*_lldA_*reporter strains and found that two alpha-hydroxy carboxylates, α-HB and glycolate, affected expression of one or both of the *lldPDE* and *lldA* loci. α-HB induced expression of both *lldPDE* and *lldA* but was a more potent inducer of *lldA*. The α-HB-dependent induction of *lldA* was also dependent on LldS, suggesting that this regulator may recognize α-HB. The notion that L-lactate-binding sites can also accommodate α-HB is consistent with the structural similarity of these two compounds and the fact that studies of two L-lactate dehydrogenase enzymes from *P. stutzeri* strains have been shown to oxidize α-HB (62, 63). Little is known about the abundance of α-HB, so the physiological significance of its effects on L-lactate dehydrogenase expression and its oxidation by these enzymes is unclear.

Glycolate had an inhibitory effect on gene expression and this effect was specific to *lldPDE*. This compound, which is also structurally similar to lactate, is found in diverse environments as it is present in large quantities in aquatic settings (64, 65) and has also been detected in chronic pressure ulcer wounds (66). In a study examining the transcriptomes of dual-species biofilms, the *P. aeruginosa glc* genes as well as *lldA* were induced during coculture with *S. aureus* when compared to monoculture conditions (67), suggesting that *S. aureus* produces both glycolate and L-lactate. In addition to these exogenous sources of glycolate, metabolic pathways within *P. aeruginosa* have the potential to produce this compound. A metabolome analysis of *P. aeruginosa* PAO1 detected enhanced glycolate levels under conditions that stimulated flux through the glyoxylate shunt, indicating that glycolate may be a byproduct of this pathway (68). Another potential source of endogenous glycolate is detoxification pathways for glyoxals, which are toxic side-products of metabolic processes and are often associated with oxidative stress (69). In organisms where glyoxal detoxification has been studied, it occurs via conversion of glyoxal into glycolate by the enzymes glyoxalase I and II. *P. aeruginosa* contains homologs to these enzymes and therefore has the potential to carry out this pathway (70). Our results, which indicate that glycolate production is localized to the biofilm-air interface of macrocolonies, are consistent with both potential sources of glycolate because this region is subject to carbon limitation (a condition that promotes use of the glyoxylate shunt) and oxidative stress (a condition that would be expected to correlate with glyoxal production and detoxification) (71–74).

In studies examining the physiological heterogeneity and architecture of *P. aeruginosa* macrocolonies, we have used stimulated Raman scattering microscopy to detect metabolic activity (75) across biofilm depth. Our results show that metabolic activity levels are spatially heterogeneous in biofilms. They also show that, in biofilms grown on complex or defined media and with a range of carbon source concentrations, maximum metabolic activity occurs at a distance from the biofilm-air interface (19, 74, 76). This suggests that the hypoxic biofilm subzone is more conducive to metabolism than the oxic region, and raises the possibility that the redundant L-iLDH enzymes are optimized to function in, and may play a role in determining, the different levels of metabolic activity occurring in biofilm subzones. Interestingly, despite the marked differences we observed in the responses of *lldD* and *lldA* to metabolic cues, we found that both loci contribute to persistence within macrophages indicating that while multicellular growth can promote differential use of LldD and LldA, an in vivo condition has the potential to promote their simultaneous use. Together, our observations provide a case study of the conditional use of redundant enzymes and provide insight into the metabolic strategies employed by a devastating bacterium that forms multicellular structures and survives intracellularly during infection (77–81).

## Methods

### Bacterial strains and culture conditions

*Pseudomonas aeruginosa* strain UCBPP-PA14 (“PA14”) was used for all experiments. Overnight cultures were grown in Lysogeny Broth (LB) (82) shaking at 200 rpm at 37°C for 16-18 hours.

### Construction of markerless deletions and fluorescent reporter strains

Approximately 1 kb of flanking sequence from each side of the target locus were amplified using the primers listed in **Table S3** and inserted into pMQ30 through gap repair cloning in *Saccharomyces cerevisiae* InvSc1. Plasmids used in this study are listed in **Table S2**. Each plasmid was transformed into *Escherichia coli* strain UQ950, verified by sequencing, and moved into *P. aeruginosa* PA14 using biparental conjugation via the *E. coli* donor strain BW29427. PA14 single recombinants were selected on LB agar plates containing 70 μg/mL gentamicin. Double recombinants (markerless mutants) were selected on a modified LB medium (containing 10% w/v sucrose and lacking NaCl) containing 1.5% agar and genotypes were confirmed by PCR. Combinatorial mutants were constructed by using single mutants as hosts for biparental conjugation as indicated in **Table S1**.

To construct reporter strains, promoter regions of varying length were amplified from the PA14 genome using primers listed in **Table S3** and inserted upstream of the coding sequence of *gfp, mScarlet,* or *luxCDABE* on their respective plasmids via ligation. Plasmids were transformed into *E. coli* UQ950 cells and verified by sequencing. Verified plasmids were introduced into PA14 using biparental conjugation with *E. coli* S17-1. Single recombinants were selected on agar plates with M9 minimal medium (47.8 mM Na_2_HPO_4_7H_2_O, 2 mM KH_2_PO_4_, 8.6 mM NaCl, 18.6 mM NH_4_Cl, 1 mM MgSO_4_, 0.1 mM CaCl_2_, 20 mM sodium citrate dihydrate, 1.5% agar) containing 70 µg/mL gentamicin or 150 µg/mL tetracycline for the luciferase reporters. The plasmid backbone was resolved out of PA14 using Flp-FRT recombination using the pFLP2 plasmid (83) and selection on M9 minimal medium 1.5% agar plates containing 300 µg/mL carbenicillin. Strains were cured of the pFLP2 plasmid by streaking on LB agar plates without NaCl and with 10% w/v sucrose. The presence of *gfp, mScarlet,* or *luxCDABE* in final clones was confirmed by PCR.

### Liquid culture growth assays

Biological triplicates of overnight pre-cultures were diluted 1:100 in 200 μL MOPS medium (50 mM MOPS, 43 mM sodium chloride, 93 mM ammonium chloride, 2.2 mM monobasic potassium phosphate, 1 mM magnesium sulfate), containing carbon sources as indicated, in a flat bottom, black polystyrene 96-well plate (Greiner Bio-One 655001). Ferrous sulfate was added, at a concentration of 1 µg/mL (3.5 µM), to the MOPS medium unless otherwise indicated. Carbon sources (succinate, lactate, glycolate, etc.) were added as sodium salts at the concentration indicated. Plates were incubated at 37°C with continuous shaking on the medium setting in a Biotek Synergy H1 plate reader. Expression of mScarlet, GFP, or luciferase was assessed by taking fluorescence or luminescence readings every 30 min for up to 24 h. mScarlet was detected at excitation and emission wavelengths of 569 nm and 599 nm, respectively. GFP was detected at excitation and emission wavelengths of 480 nm and 510 nm, respectively. Growth was assessed by taking OD readings at 500 nm simultaneously with the fluorescence/luminescence readings. Unless otherwise stated, the reported values were obtained by first subtracting the fluorescence values of “no-reporter” strains and then normalizing to growth (OD at 500 nm).

### Biolog metabolite screens

Phenotypic microarray plate PM1 (Biolog, Cat. No. 12111) contains a unique carbon source in each well of a 96-well plate (**Supplemental File 2**). For activator screens, 100 µL of MOPS medium containing 20 mM sodium succinate was added to each well of this phenotypic microarray plate, and for inhibitor screens, 100 µL of MOPS medium containing 20 mM sodium succinate and 5 mM sodium L-lactate was added to each well. Plates were incubated shaking at 37°C for approximately one hour to fully dissolve each compound. The compound solutions were then transferred to a black 96-well plate pre-filled with 100 µL of the base medium per well, so that the total volume per well was 200 µL. Overnight pre-cultures were diluted 1:100 into each well. Plates were placed in a Biotek Synergy H1 plate reader and experimental parameters were as described above. A compound was considered a “hit” if the raw fluorescence value in its well (at 7.5 hours of incubation; representing early stationary phase) was two standard deviations above (for activators) or below (for inhibitors) the median raw fluorescence value for all wells. Each screen was performed in two independent experiments, and only metabolites that were identified as hits in both experiments are reported.

### Titration experiments with luciferase reporters

The same protocol was used as for liquid culture growth assays, but with different concentrations of L-lactate, α-HB, or ferrous sulfate used to supplement the MOPS base medium. The L-lactate and α-HB titrations were performed in MOPS medium containing 20 mM succinate with L-lactate concentrations ranging from 0.1 µM to 30 mM and α-HB concentrations from 10 µM to 30 mM. The ferrous sulfate titration was done in MOPS containing 40 mM L-lactate with the concentration of added ferrous sulfate ranging from 5 nM to 1 µM. Plates were placed in a Biotek Synergy H1 plate reader and experimental parameters were the same as described above. Cultures were grown for 15-24 hours and the maximum luminescence value was identified for each concentration, normalized to growth and to the luminescence values recorded in a promoterless *lux* strain. For lactate and α-HB titrations, this value was also normalized by subtracting the maximum fluorescence value recorded during the full incubation period in the base medium (i.e., without added L-lactate/α-HB). Where indicated, the maximum luminescence value for each L-lactate/α-HB concentration is expressed as its proportion of the maximum value exhibited for that strain at any concentration. For P*_lldP_-lux*, the maximum luminescence value was detected during growth in 500 µM L-lactate, while for P*_lldA_-lux*, the maximum luminescence value was detected during growth in 30 mM L-lactate or 30 mM α-HB (the highest concentrations tested).

### AlphaFold calculations

AlphaFold-Multimer (84) was used to predict models of an LldS dimer, which were manually examined with PyMOL and Coot (85). For the model of the L-lactate complex, a search for structural homologs of LldS in the Protein Data Bank (PDB) was carried out with the program Dali (86). The structure of AmpR effector binding domain in complex with a peptide ligand (PDB entry 4WKM) (28) was found to be a close homolog, and the C-terminal Ala residue of the peptide ligand was used as the starting point for modeling L-lactate. The hydrogen-bonding interactions between the carboxylate group of Ala and the protein was maintained for L-lactate, while the position of the rest of the compound was adjusted manually. The Thr and Tyr residues in AmpR that are hydrogen-bonded to the C-terminal carboxylate of the peptide are conserved in LldS, and these residues have the same conformation in the two structures.

### Protein expression and purification

PA14 *lldS* was cloned into the pET28a vector with a N-terminal 6xHis-tag. The proteins were overexpressed in *E. coli* BL21 (DE3) cells at 16°C overnight. For purification, the cell pellet was resuspended and lysed by sonication in a buffer containing 20 mM Tris (pH 8.0), 500 mM NaCl, 2 mM bME, 5% (v/v) glycerol, and one tablet of the protease inhibitor mixture (Sigma). The cell lysate was then centrifuged at 13,000 rpm for 30 min at 4°C. The protein was purified from the supernatant via nickel affinity chromatography (Qiagen). The protein was further purified by a Hiload 16/60 Superdex 200 column (Cytiva) equilibrated with a buffer containing 20 mM Tris (pH 8.0), 500 mM NaCl, and 2 mM DTT. The purified protein was concentrated to 20 mg/ml, supplemented with 5% (v/v) glycerol, and stored at -80°C.

### Fluorescence anisotropy

Fluorophore-labeled DNA was generated using PCR amplification with a 5’FAM-labeled reverse primer (IDT) and an unlabelled forward primer and purified using the E.Z.N.A Gel Extraction Kit (Omega Bio-tek). The amplified region contained the 256-bp region directly upstream of the *lldA* start codon. Purified LldS protein and 5 nM of FAM-labeled probe were added to reaction buffer (50 mM HEPES (pH 7.5), 30 mM KCl, 3 mM magnesium acetate, 5% glycerol) with protein concentration ranging from 9.37 nM to 5 µM. Parallel and perpendicular polarization values were measured in a black 384-well plate using a Biotek Synergy Neo2 plate reader with excitation and emission wavelengths of 485 nm and 528 nm, respectively and fluorescence anisotropy was calculated using the formula: (Parallel - Perpendicular)/Parallel + (2 * Perpendicular) (87). Before plotting, the minimum anisotropy value was subtracted from all data points. K_d_ was calculated in GraphPad Prism by curve-fitting using a non-linear, least squares regression model and assuming one-site, specific binding.

### Gene arrangement and association analysis

Genomic sequences (*i.e.* GenBank files) of 1,323 strains were downloaded from the Pseudomonas DB (https://pseudomonas.com/) (34). For *P. aeruginosa*, only one strain, PA14, was chosen, while strains from all other species were kept for analysis. This resulted in 689 strains for the search of lactate dehydrogenase-related genes. The gene arrangement search was implemented in a custom-built tool named “Locus Hunter” (https://github.com/linyc74/locus_hunter). For each gene arrangement, blastp was used to search for homologous genes (e-value = 10^-60^) (88). The interval of each homologous gene was extended by a flanking length of 5,000 bp, resulting in intervals that overlap with each other. Intervals that overlap with each other were then fused to form a gene arrangement containing genes of interest. For each species, only one representative strain was chosen; however, if two or more strains showed different gene arrangements both were represented. These considerations reduced the number of strains to 213. For the association analysis, two gene arrangements were defined to be associated when they coexist in a given strain. The number of associations between a pair of gene arrangements is the number of strains harboring both arrangements.

### Phylogenetic analysis

Out of the 213 strains identifed in the gene arrangement analysis (described above), 179 strains contained *lldD* and *lldA* homologs. In Geneious Prime (Version 2024.0.3), the corresponding LldD and LldA protein sequences were aligned using Clustal Omega and a tree was constructed using Geneious Tree Builder (Jukes-Cantor; Neighbor-Joining; No Outgroup).

### Preparation of *P. aeruginosa* macrocolony biofilms

Overnight *P. aeruginosa* cultures were subcultured 1:100 in 3 mL LB and were grown at 37°C, 200 rpm to an OD500 of ∼0.5-0.6. Then, 5 µL of liquid subcultures were spotted onto 1% agar plates (100 mm x 15 mm, Simport Scientific D210-16) containing 60 mL of MOPS medium with 20 mM sodium succinate and 10 mM sodium L-lactate. For biofilms used for thin-sectioning experiments, agar plates were prepared in two layers: a 45 mL base layer and a 15 mL top layer. The base MOPS medium was prepared the same in every case except in experiments testing the effect of iron availability, in which ferrous sulfate was omitted for the “0 µM” condition and added in excess for the “10 µM” condition. For mixed biofilms containing two reporter strains, the OD500 of each subculture was corrected to exactly 0.5 and subcultures were mixed in a 1:1 ratio before being spotted. Macrocolony biofilms were grown in the dark at 25°C. Macrocolony biofilm experiments were conducted with at least three biological replicates of each strain and condition.

### Top-down fluorescence imaging of *P. aeruginosa* macrocolony biofilms

After 3 days of growth as described above, bright-field and fluorescence images were obtained using a Zeiss Axio Zoom V16 fluorescence stereo zoom microscope (excitation, 545 nm; emission, 605 nm for imaging of mScarlet). Images were processed using the Zeiss Zen software and analyzed using Fiji/ImageJ.

### Quantification of colony forming units (CFUs) from macrocolony biofilms

After 3 days of growth as described above, biofilms were homogenized in 1 mL PBS using a bead mill homogenizer (Omni [Kennesaw, GA] Bead Ruptor 12; at high setting for 90 s) and 0.5 g ceramic beads (Thermo Fisher 15 340 159, diameter of 1.4 mm). The cell suspension was serially diluted in PBS, plated on 1% tryptone, 1.5% agar plates and incubated at 37°C for 16 hours before CFU counting.

### Thin sectioning of PA14 macrocolony biofilms

After 3 days of growth on bilayer agar plates as described above, biofilms were overlaid with 15 mL 1% agar and sandwiched biofilms were lifted from the bottom agar layer and fixed overnight in 4% paraformaldehyde in PBS at room temperature for 24 hours. Fixed biofilms were processed, infiltrated with wax, sectioned in 10-um-thick sections, and collected onto slides as described in (76). Slides were air-dried overnight, heat-fixed on a hotplate for 30 minutes at 45°C, and rehydrated in the reverse order of processing. Rehydrated colonies were immediately mounted in TRIS-buffered DAPI:Fluorogel (Electron Microscopy Sciences) and overlaid with a coverslip. Thin sections were imaged using a Zeiss Axio Zoom V16 fluorescence stereo zoom microscope, at the same settings as described above. Images were processed using the Zeiss Zen software and analyzed using Fiji/ImageJ.

### Assessment of intracellular bacterial load in macrophages

RAW264.7 macrophage cells were plated overnight on six-well plates. Bacterial strains (WT, Δ*lldD*, Δ*lldA*, Δ*lldS*, and Δ*lldD*Δ*lldA*) were grown in LB to an OD600 of 2.0 and then washed with PBS. Macrophages were washed with antibiotic-free Dulbecco’s Modified Eagle Medium (DMEM) and infected with bacteria at 5 MOI for 30 min at 37°C in 5% CO_2_. Bacteria that were unbound to macrophages were removed by one wash with cold DMEM medium, and to remove any excess extracellular bacteria, 100 µg/ml of gentamicin was added for 30 min; cells were washed with DMEM and transferred to medium without gentamicin and incubated at 37°C in 5% CO_2_ for 3 hours. Infected RAW cells were lysed in distilled water. The lysed cells were immediately diluted in PBS and plated on LB agar plates to assess the intracellular bacterial load in macrophages. Bacterial CFU were counted after incubating the plates overnight at 37°C.

### Fluorescent imaging of macrophages

RAW264.7 cells were seeded at 1×10^5^ cells per well on three-well-chambered coverglass slides the day prior to the experiment. Cells were infected with WT PA14 or PA14 bacteria carrying the P*_lldA_*-*mScarlet* reporter at an MOI of 5, as described in the previous section. Cells were then fixed in 4% paraformaldehyde in PBS (pH 7.4) for 15 min at room temperature. Slides were washed in 1 x PBS (pH 7.4). Cells were stained with DAPI (1:10,000) at room temperature and washed in 1 x PBS (pH 7.4) three times for 5 min each. The cells were examined using a confocal microscope (Nikon ECLIPSE Ti2; Nikon Instruments Inc., Tokyo) at 400x optical magnification. The assay was conducted in triplicate and repeated three times.

## Acknowledgments

This work was supported by NIH/NIAID grant R01AI103369 to L.E.P.D., R35GM118093 to L.T. and NIH awards R01AI134857, R01AI177555, and Shriner’s grant 83009 to L.G.R. The authors thank Hannah Peha for assistance with strain engineering and Riley Gentry, Nicholas Ide, Ji Huang, and Columbia’s Precision Biomolecular Characterization Facility (PBCF) for assistance with the anisotropy experiments.

