## Supplemental figure S1-7 for "The L-lactate dehydrogenases of *Pseudomonas aeruginosa* are conditionally regulated but both contribute to survival during macrophage infection"

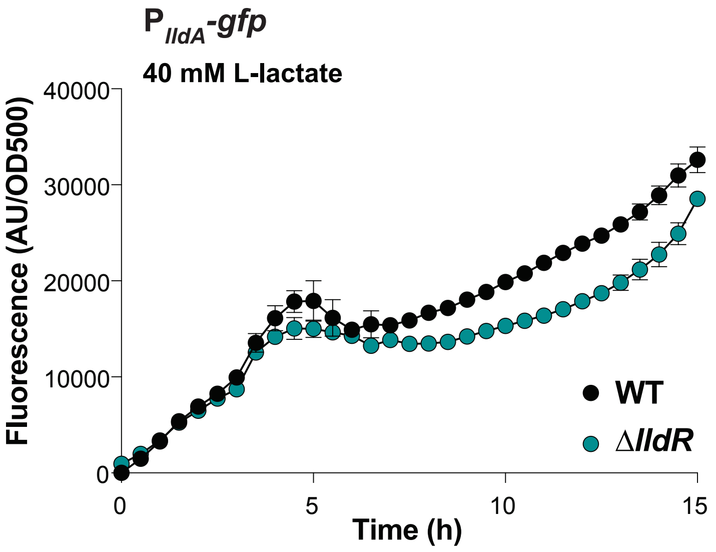


**Supplemental Figure 1. LldR is not a major effector of *lldA* expression.** *lldA* promoter activity in liquid cultures of WT and Δ*lldR* grown in MOPS medium containing 40 mM L-lactate.

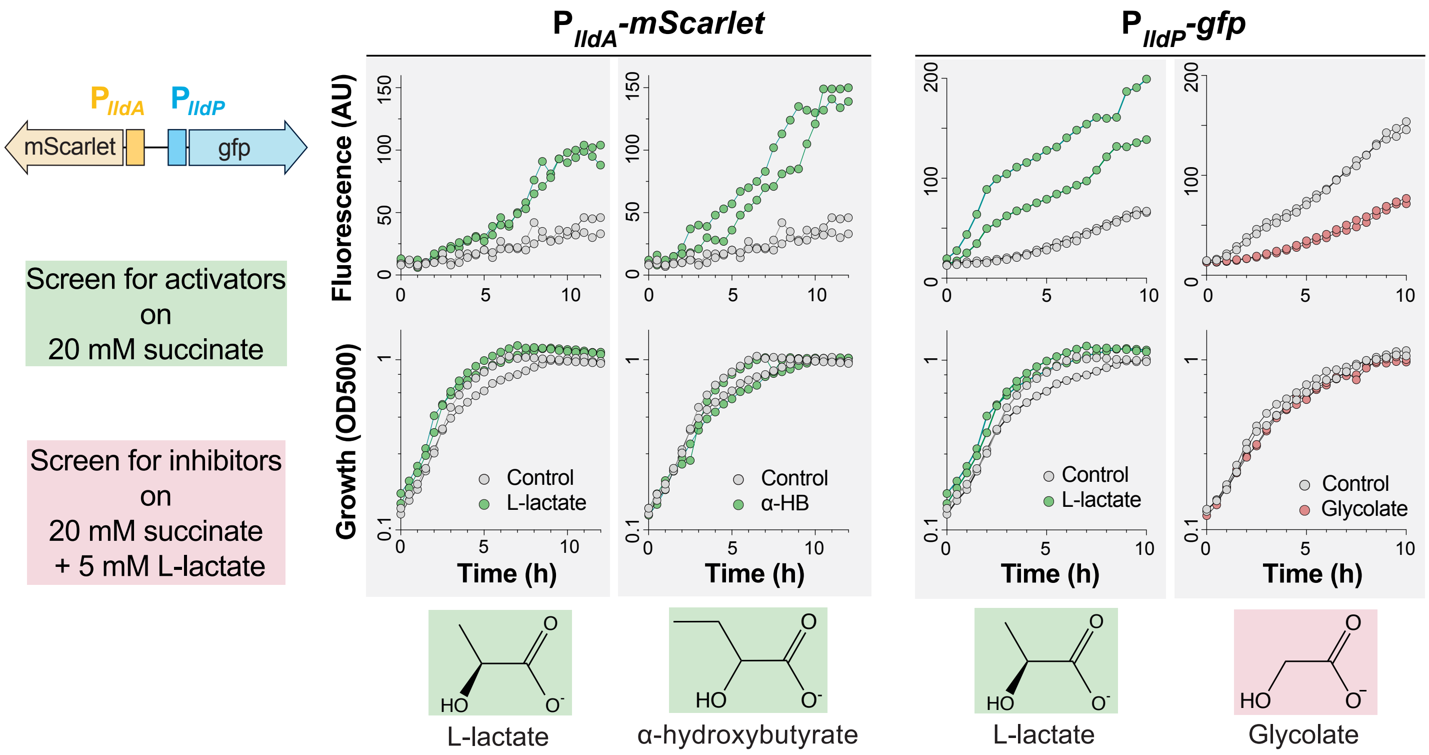


**Supplemental Figure 2. Activators and inhibitors of *lldA* and *lldPDE* expression identified by screening with plate PM-1 (Biolog, Inc.)**. Left: Schematic of constructs in the dual-reporter strain used in the screens. Conditions for activator and inhibitor screens are shown. Center and right: Raw fluorescence curves, growth curves, and chemical structures of compounds identified in the screen for small molecules affecting *lldA* (center) or *lldP* (right) promoter activity. Two lines are shown per condition, representing two independent experiments. Results for activators are plotted in green, those for inhibitors are plotted in red, and those for control wells are plotted in gray.

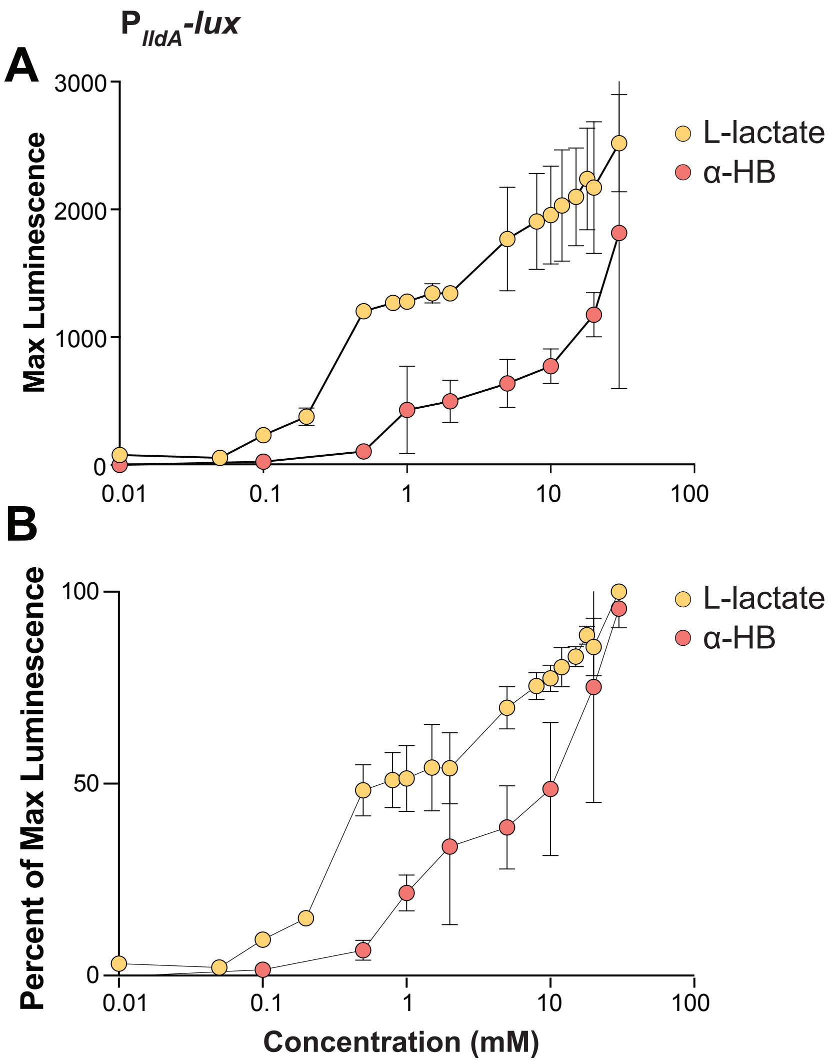


**Supplemental Figure 3. Effect of various α-HB concentrations on *lldA* expression**. Activity of the *lldA* promoter at α-HB concentrations ranging from 10 μM to 30 mM (pink data points), with the L-lactate titration from Figure 1C provided for comparison (yellow data points). Cultures of the P$_{lldA}$-*lux* reporter strain were grown shaking in a 96-well plate at 37°C for 24 hours in a base medium of MOPS containing 20 mM succinate. (A) Each value shown represents the maximum luminescence produced during growth in the indicated condition. (B) Values from A were normalized to the maximum luminescence value produced in the most-stimulatory concentration. Values shown for each concentration are averages of two biological replicates and error bars represent standard deviation.

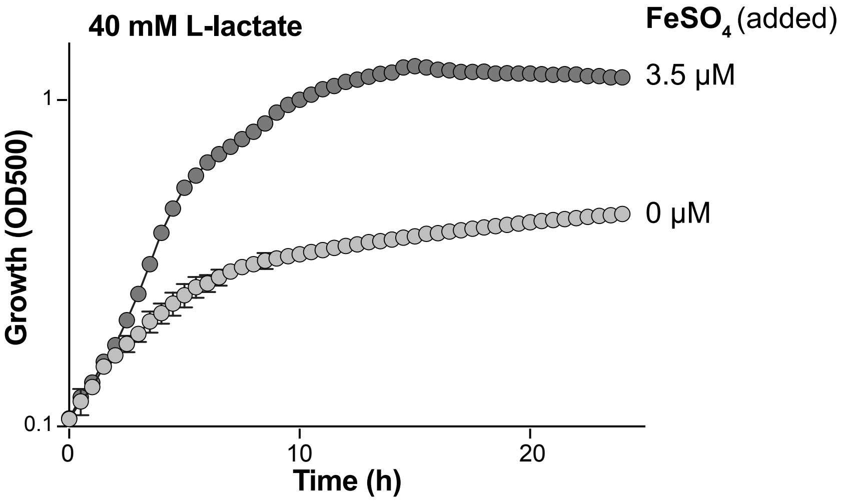


**Supplemental Figure 4. Liquid culture growth is limited in MOPS medium without added ferrous sulfate.** Growth (optical density at 500 nm) of WT in liquid MOPS medium containing 40 mM L-lactate and either 0 or 3.5 µM of added iron.

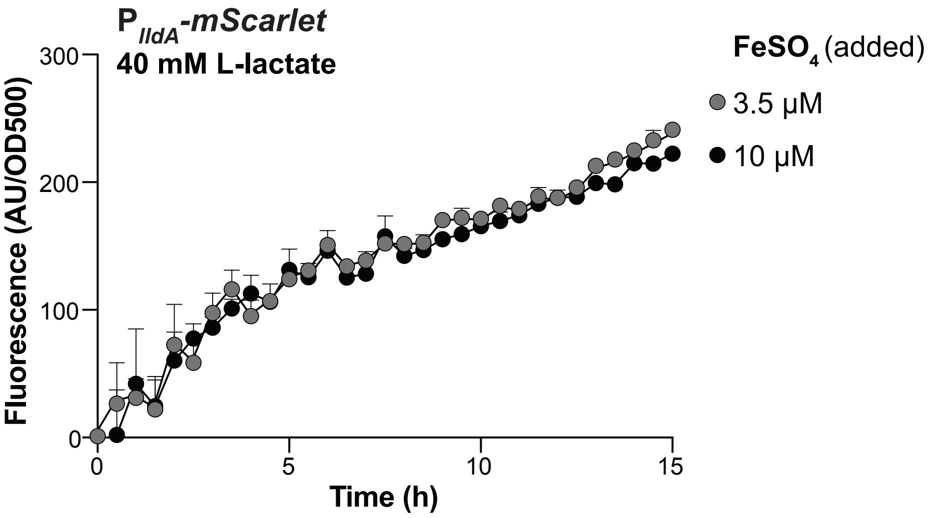


**Supplemental Figure 5. *lldA* expression is unaffected by the addition of increased ferrous sulfate to the medium.** *lldA* promoter activity in liquid cultures of WT grown in MOPS medium containing 40 mM L-lactate and either 3.5 or 10 μM of added iron.

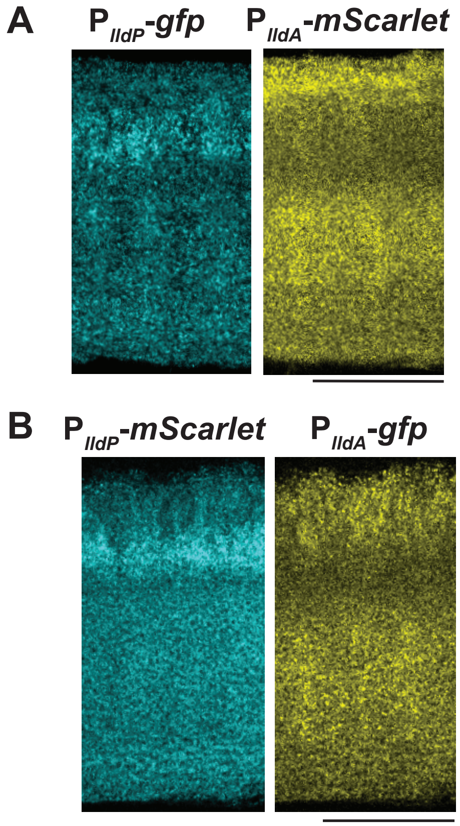


**Supplemental Figure 6. Spatial patterning of fluorescence in biofilms of the dual-reporter strain is recapitulated in mixed biofilms containing single-reporter strains.** (**A**) Fluorescence images of a thin-section from a biofilm inoculated using an equal mixture of the $P_{lldP}$-*gfp* and $P_{lldA}$-*mScarlet* reporter strains. *mScarlet* fluorescence is shown in yellow and *gfp* fluorescence is shown in cyan. (**B**) Fluorescence images of a thin-section from a biofilm inoculated using an equal mixture of $P_{lldP}$-*mScarlet* and $P_{lldA}$-*gfp* reporter strains. *mScarlet* fluorescence is shown in cyan and *gfp* fluorescence is shown in yellow. Biofilms were grown on MOPS medium containing 20 mM succinate and 10 mM L-lactate.

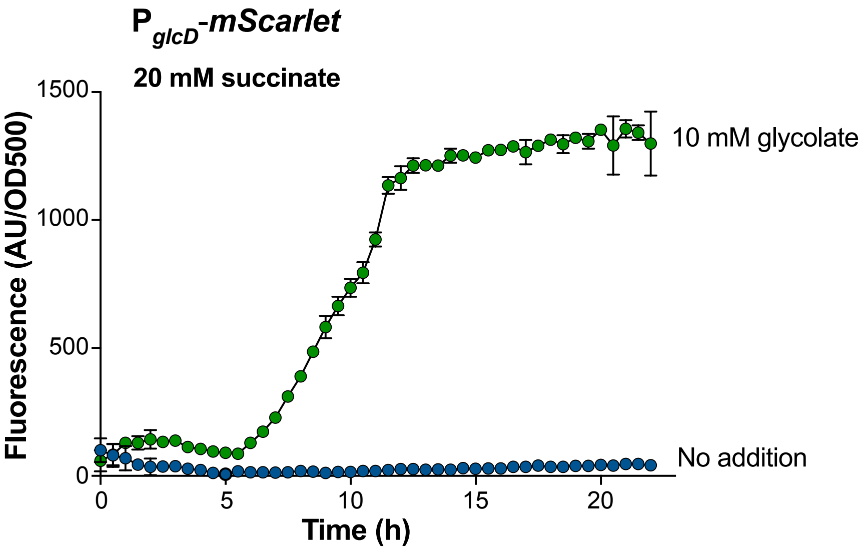


**Supplemental Figure 7. Expression of the *glcDEFG* operon is induced by added glycolate.** *glcD* promoter activity in liquid cultures of WT grown in MOPS medium containing 20 mM succinate (blue data points) or the same medium with glycolate added at 10 mM (green data points).
