## Supplemental tables S1-3 for "The L-lactate dehydrogenases of *Pseudomonas aeruginosa* are conditionally regulated but both contribute to survival during macrophage infection"

**Table S1. Bacterial strains used in this study.**

| Number | Strain | Description | Source |
|---|---|---|---|
| ***Pseudomonas aeruginosa* strains** | | | |
| LD0 | UCBPP-PA14 (WT) | Clinical isolate UCBPP-PA14. | (1) |
| LD2798 | PA14 *attB*::$P_{lldP}$-*gfp* | PA14 containing a construct in the *attB* site that expresses *gfp* under control of the 324 bp region upstream of *lldP* (*PA14_63080*). Made using pLD2797. | (2) |
| LD2868 | PA14 *attB*::$P_{lldA}$-*gfp* | PA14 containing a construct in the *attB* site that expresses *gfp* under control of the 356 bp region upstream of *lldA* (*PA14_33860*). Made using pLD2867. | (2) |
| LD4472 | PA14 *attB*::$P_{lldP}$-*lux* | PA14 containing a construct in the *attB* site that expresses the *luxCDABE* operon under control of the 324 bp region upstream of *lldP* (*PA14_63080*). | This study |
| LD4768 | PA14 *attB*::$P_{lldA}$-*lux* | PA14 containing a construct in the *attB* site that expresses the *luxCDABE* operon under control of the 356 bp region upstream of *lldA* (*PA14_33860*). | This study |
| LD4486 | PA14 *attB*::MCS-*lux* | PA14 containing a construct in the *attB* site that expresses the *luxCDABE* operon with no upstream promoter. Made using pLD4440. | This study |
| LD4157 | PA14 *attB*::$P_{lldP}$-*mScarlet* | PA14 containing a construct in the *attB* site that expresses *mScarlet* under control of the 324 bp region upstream of *lldP* (*PA14_63080*). Made using pLD4111. | This study |
| LD3760 | PA14 *attB*::$P_{lldA}$-*mScarlet* | PA14 containing a construct in the *attB* site that expresses *mScarlet* under control of the 356 bp region upstream of *lldA* (*PA14_33860*). Made using pLD3738. | This study |
| LD4189 | PA14 *attB*::$P_{lldA}$-*mScarlet,* $P_{lldP}$-*gfp* | PA14 containing a dual reporter construct in the *attB* site that expresses *mScarlet* under control of the 356 bp region upstream of *lldA* (*PA14_33860*) and *gfp* under control of the 324 bp region upstream of *lldP* (*PA14_63080*). Made using pLD4165. | This study |
| LD4187 | Δ*lldD* | PA14 with *lldD* (*PA14_63090*) deleted. Made by mating pLD4132 into UCBPP-PA14. | This study |
| LD4052 | Δ*lldA* | PA14 with *lldA* (*PA14_33860*) deleted. Made by mating pLD2758 into UCBPP-PA14. | (2) |
| LD3714 | Δ*lldS* | PA14 with *lldS* (*PA14_33840*) deleted. Made by mating pLD3690 into UCBPP-PA14. | This study |
| LD4862 | Δ*lldD* Δ*lldA* | PA14 with *lldD* (*PA14_63090*) and *lldA* (*PA14_33860*) deleted. Made by mating pLD2758 into LD4187. | This study |
| LD5130 | Δ*lldD* Δ*lldS* | PA14 with *lldD* (*PA14_63090*) and *lldS* (*PA14_33840*) | This study |

| | | | |
|---|---|---|---|
| | | deleted. Made by mating pLD4132 into LD3714. | |
| LD2735 | Δ*lldD* Δ*lldE* | PA14 with *lldD* (*PA14_63090*) and *lldE* (*PA14_63100*) deleted. Made by mating pLD2734 into UCBPP-PA14. | (2) |
| LD3715 | Δ*lldS attB*::$P_{lldA}$-*gfp* | Δ*lldS* containing a construct in the *attB* site that expresses *gfp* under control of the 356 bp region upstream of *lldA* (*PA14_33860*). Made by mating pLD3690 into LD2868. | This study |
| LD5116 | Δ*lldS* *attB*::$P_{lldA}$-*mScarlet* | Δ*lldS* containing a construct in the *attB* site that expresses *mScarlet* under control of the 356 bp region upstream of *lldA* (*PA14_33860*). Made by mating pLD3690 into LD3760. | This study |
| LD3809 | Δ*lldS*::*lldS* | Δ*lldS* (*PA14_33840*) strain with wild-type *lldS* complemented back into the site of deletion. Made by mating pLD3797 into LD3714. | This study |
| LD3810 | Δ*lldS*::*lldS* *attB*::$P_{lldA}$-*gfp* | Δ*lldS* (*PA14_33840*) containing a construct in the attB site that expresses gfp under the control of the 356 bp region upstream of *lldA* with wild-type *lldS* complemented back into the site of deletion. Made by mating pLD3797 into LD3715. | This study |
| LD5161 | Δ*lldS*::$LldS^{T107M}$ | T107 of *lldS* (*PA14_33840*) mutated to methionine in the native locus. Made by mating pLD5150 into LD3714. | This study |
| LD5162 | Δ*lldS*::$LldS^{T107M}$ *attB*::$P_{lldA}$-*gfp* | T107 of *lldS* (*PA14_33840*) mutated to methionine in the native locus. Containing a construct in the attB site that expresses gfp under the control of the 356 bp region upstream of *lldA*. Made by mating pLD5150 into LD3715. | This study |
| LD5169 | Δ*lldS*::$LldS^{T107A}$ | T107 of *lldS* (*PA14_33840*) mutated to alanine in the native locus. Made by mating pLD5173 into LD3714. | This study |
| LD5170 | Δ*lldS*::$LldS^{T107A}$ *attB*::$P_{lldA}$-*gfp* | T107 of *lldS* (*PA14_33840*) mutated to alanine in the native locus. Containing a construct in the attB site that expresses gfp under the control of the 356 bp region upstream of *lldA*. Made by mating pLD5173 into LD3715. | This study |
| LD3508 | Δ*lldR* | PA14 with *lldR* (*PA14_63070*) deleted. Made by mating pLD3512 into UCBPP-PA14. | This study |
| LD3516 | Δ*lldR attB*::$P_{lldP}$-*gfp* | Δ*lldR* (*PA14_63070*) containing a construct in the *attB* site that expresses *gfp* under control of the 324 bp region upstream of *lldP* (*PA14_63080*). Made by mating pLD3512 into LD2798. | This study |
| LD3510 | Δ*lldR attB*::$P_{lldA}$-*gfp* | Δ*lldR* (*PA14_63070*) containing a construct in the *attB* site that expresses *gfp* under control of the 356 bp region upstream of *lldA*. Made by mating pLD3512 into | This study |

| | | | |
|---|---|---|---|
| | | LD2868. | |
| LD4654 | Δ*fur2* | PA14 with *fur2* (*PA14_33830*) deleted. Made by mating pLD4660 into UCBPP-PA14. | This study |
| LD4655 | Δ*fur2* *attB*::$P_{lldA}$-*mScarlet* | Δ*fur2* containing a construct in the *attB* site that expresses *mScarlet* under control of the 356 bp region upstream of *lldA* (*PA14_33860*). Made by mating pLD4660 into LD3760. | This study |
| LD4394 | Δ*pvdS* | PA14 with *pvdS* (PA14_33260) deleted. | S. Häussler, Helmholtz |
| LD4557 | Δ*pvdS* *attB*::$P_{lldA}$-*mScarlet* | Δ*pvdS* containing a construct in the *attB* site that expresses *mScarlet* under control of the 356 bp region upstream of *lldA* (*PA14_33860*). Made by mating pLD3738 into LD4394. | This study |
| LD3788 | PA14 *attB*::$P_{fur2}$-*gfp* | PA14 containing a construct in the *attB* site that expresses *gfp* under control of the 390 bp region upstream of *fur2* (*PA14_33830*). Made using pLD3777. | This study |
| LD4953 | PA14 *attB*::$P_{glcD}$-*mScarlet* | PA14 containing a construct in the *attB* site that expresses *mScarlet* under control of the 494 bp region upstream of *glcD* (*PA14_70690*). Made using pLD4757. | This study |
| LD4233 | Δ*lldD* *attB*::$P_{lldA}$-*mScarlet,* $P_{lldP}$-*gfp* | Δ*lldD* containing a dual reporter construct in the *attB* site that expresses *mScarlet* under control of the 356 bp region upstream of *lldA* (*PA14_33860*) and *gfp* under control of the 324 bp region upstream of *lldP* (*PA14_63080*). Made by mating pLD4132 into LD4189. | This study |
| LD4235 | Δ*lldA* *attB*::$P_{lldA}$-*mScarlet,* $P_{lldP}$-*gfp* | Δ*lldA* containing a dual reporter construct in the *attB* site that expresses *mScarlet* under control of the 356 bp region upstream of *lldA* (*PA14_33860*) and *gfp* under control of the 324 bp region upstream of *lldP* (*PA14_63080*). Made by mating pLD2758 into LD4189. | This study |
| LD5274 | PA14 *attTn7*::$P_{PA1/04/03}$-*mScarlet* | PA14 containing a construct in the attTn7 site that expresses the coding region of *mScarlet* under control of the lac-derived constitutive *PA1/04/03* promoter. | This study |
| LD4600 | PA14 *attB*::$P_{lldA}$*(256)-gfp* | PA14 containing a construct in the *attB* site that expresses *gfp* under control of the 256 bp region upstream of *lldA*. Made using pLD4600. | This study |
| LD4980 | PA14 *attB*::$P_{lldA}$*(221)-gfp* | PA14 containing a construct in the *attB* site that expresses *gfp* under control of the 221 bp region upstream of *lldA*. Made using pLD4966. | This study |
| LD4982 | PA14 *attB*::$P_{lldA}$*(188)-gfp* | PA14 containing a construct in the *attB* site that expresses *gfp* under control of the 188 bp region upstream of *lldA*. Made using pLD4967. | This study |

| | | | |
|---|---|---|---|
| LD5027 | PA14 *attB*::$P_{lldA}$*(164)-gfp* | PA14 containing a construct in the *attB* site that expresses *gfp* under control of the 164 bp region upstream of *lldA*. Made using pLD5029. | This study |
| LD4599 | PA14 *attB*::$P_{lldA}$*(125)-gfp* | PA14 containing a construct in the *attB* site that expresses *gfp* under control of the 125 bp region upstream of *lldA*. Made using pLD4599. | This study |
| LD4598 | PA14 *attB*::$P_{lldA}$*(105)-gfp* | PA14 containing a construct in the *attB* site that expresses *gfp* under control of the 105 bp region upstream of *lldA*. Made using pLD4598. | This study |
| ***E. coli* strains** | | | |
| LD44 | UQ950 E. coli | DH5α λ(*pir*) strain for cloning; F-Δ(*argF-lac*)169Φ80d*lacZ*58 (ΔM15) *glnV44*(AS) *rfbD1 gyrA96*(NalR) *recA1 endA1 spoT thi-1 hsdR17 deoR λpir*+ D | D. Lies, Caltech |
| LD661 | BW29427 | Donor strain for conjugation: *thrB1004 pro thi rpsL hsdS lacZ* ΔM15RP4–1360 Δ(*araBAD*)*567* Δ*dapA1341*::[*erm pir*(wt)] | W. Metcalf, University of Illinois |
| LD69 | β2155 | Helper strain. *thrB1004 pro thi strA hsdsS lac*ZΔM15 (F' *lacZ*ΔM15 $lacI^q$ *traD36* $proA^+$ $proB^+$ ) Δ*dapA*::*erm* ($Erm^R$) *pir*::RP4 [::kan ($Km^R$) from SM10] | (3) |
| LD2901 | S17-1 | $Str^R$ , $Tp^R$ , F-RP4-2-*Tc*::*Mu aphA*::*Tn7 recA λpir* lysogen | (4) |
| | BL21 (DE3) | Competent *E. coli* strain used for protein purification. | New England BioLabs |
| ***Saccharomyces cerevisiae* strains** | | | |
| LD676 | InvSc1 | *MATα/MATα leu2/leu2 trp1-289/trp1-289 ura3-52/ura3-52 his3-Δ1/his3-Δ1* | Invitrogen |

**Table S2. Plasmids used in this study.**

| Plasmid name | Description | Source |
|---|---|---|
| pMQ30 | Yeast-based allelic-exchange vector; *sacB*+, CEN/ARSH, URA3+, $Gm^R$. | (5) |
| pFLP2 | Site-specific excision vector with cI857-controlled FLP recombinase. encoding sequence, *sacB*+ , $Amp^R$ . Used to insert LD2722-based plasmids into *P. aeruginosa* strains. | (6) |
| pLD2722 | mini-CTX derived plasmid. $Gm^R$ , $Tet^R$ flanked by Flp recombinase target (FRT) sites to resolve out resistance cassettes. For making GFP reporters. | (7) |
| pLD3208 | mini-CTX derived plasmid. $Gm^R$ , $Tet^R$ flanked by Flp recombinase target (FRT) sites to resolve out resistance cassettes. For making mScarlet reporters. | (8) |
| pLD4440 | pMINI-lux, a mini-CTX derivative containing promoterless *luxCDABE* operon, $Tet^R$ | (9) |
| pLD3433 | mini-Tn7 derived plasmid. $Gm^R$ , $Cm^R$ $P_{PA1/04/03}$::*mScarlet* | (10) |
| pLD2797 | 324 bp upstream of *lldP* PCR fragment in pLD2722. | (2) |
| pLD2867 | 356 bp upstream of *lldA* PCR fragment in pLD2722. | (2) |
| pLD4111 | 800 bp *mScarlet-terminator* PCR fragment ligated into pLD2797 using SacI and XhoI. | This study |
| pLD3738 | 800 bp *mScarlet-terminator* PCR fragment ligated into pLD2867 using SacI and XhoI. | This study |
| pLD4441 | 324 bp upstream of *lldP* PCR fragment ligated into pLD4440 using XhoI and EcoRI. | This study |
| pLD4769 | 356 bp upstream of *lldA* PCR fragment ligated into pLD4440 using XhoI and EcoRI. | This study |
| pLD4165 | 2943 bp PCR fragment amplified from pLD3738 containing the 356 bp *lldA* promoter and *mScarlet-terminator* sequence ligated into pLD2797 using SpeI and BamHI. | This study |
| pLD3777 | 390 bp upstream of *fur2* (*PA14_33830*) PCR fragment ligated into pLD2797 using XhoI. | This study |
| pLD4757 | 494 bp upstream of *glcD* (*PA14_70690*) PCR fragment ligated into pLD3208 using SpeI and EcoRI. | This study |
| pLD4600 | 256 bp upstream of *lldA* (*PA14_33860*) PCR fragment ligated into pLD2722 using SpeI and XhoI. | This study |
| pLD4966 | 221 bp upstream of *lldA* (*PA14_33860*) PCR fragment ligated into pLD2722 using SpeI and XhoI. | This study |

| | | |
|---|---|---|
| pLD4967 | 188 bp upstream of *lldA* (*PA14_33860*) PCR fragment ligated into pLD2722 using SpeI and XhoI. | This study |
| pLD5029 | 164 bp upstream of *lldA* (*PA14_33860*) PCR fragment ligated into pLD2722 using SpeI and XhoI. | This study |
| pLD4599 | 125 bp upstream of *lldA* (*PA14_33860*) PCR fragment ligated into pLD2722 using SpeI and XhoI. | This study |
| pLD4598 | 105 bp upstream of *lldA* (*PA14_33860*) PCR fragment ligated into pLD2722 using SpeI and XhoI. | This study |
| pLD2758 | Δ*lldA* (*PA14_33860*) flanking fragments introduced into pMQ30 by gap repair cloning in yeast strain InvSc1. | (2) |
| pLD2734 | Δ*lldDE* (*PA14_63090-63100*) flanking fragments introduced into pMQ30 by gap repair cloning in yeast strain InvSc1. | (2) |
| pLD4132 | Δ*lldD* (*PA14_63090*) flanking fragments introduced into pMQ30 by gap repair cloning in yeast strain InvSc1. | This study |
| pLD3690 | Δ*lldS* (*PA14_33840*) flanking fragments introduced into pMQ30 by gap repair cloning in yeast strain InvSc1. | This study |
| pLD4660 | Δ*fur2* (*PA14_33830*) flanking fragments introduced into pMQ30 by gap repair cloning in yeast strain InvSc1. | This study |
| pLD3512 | Δ*lldR* (*PA14_63070*) flanking fragments introduced into pMQ30 by gap repair cloning in yeast strain InvSc1. | This study |
| pLD3797 | The CDS of *lldS* with ~1 kb flanks on either side, introduced into pMQ30 by gap repair cloning in yeast strain InvSc1. | This study |
| pLD5150 | The CDS of *lldS* with ~1 kb flanks on either side with T107 changed to an methionine, introduced into pMQ30 by gap repair cloning in yeast strain InvSc-1. | This study |
| pLD5173 | The CDS of *lldS* with ~1 kb flanks on either side with T107 changed to an alanine, introduced into pMQ30 by gap repair cloning in yeast strain InvSc-1. | This study |
| pET28a | Expression plasmid used for protein purification. Contains a T7 promoter, 6x-His tag. $Kan^R$ | Addgene |
| pET28a-lldS | Expression plasmid containing a T7 promoter in front of the *lldS* sequence. Contains an N-termin 6x-His tag. $Kan^R$ | This study |

**Table S3. Primers used in this study.**

| Primer Number | Sequence |
|---|---|
| Primers for plasmids pLD4111, pLD3738 (used to make $P_{lldP}$-*mScarlet,* $P_{lldA}$-*mScarlet*) | |
| 2609 | acgtacgt ctcgag tctagatttaaga aggaga tatacat ATGAGTAAAGGAGAAGC |
| 2635 | actgactg gagctc ATAAAACGAAAGGCCCAGTCTTTCG |
| Primers for plasmid pLD4441 (used to make $P_{lldP}$-*lux*) | |
| 3895 | tatctcgaggctCGACACCCTTACCCGAAGTT |
| 3896 | tatgaattcgttGGGTTGGCTCCCTAATTGTTG |
| Primers for plasmid pLD4769 (used to make $P_{lldA}$-*lux*) | |
| 4121 | tatctcgaggctTGCTCGATTTGGGCATGAC |
| 4122 | tatgaattcgttGCAGTCCACTCCTTCGGG |
| Primers for plasmid pLD4165 (used to make dual reporter $P_{lldP}$-*gfp*, $P_{lldA}$-*mScarlet*) | |
| 3781 | catagactagtctatgcgcggtcgatgcgatgtcgatagcTGCTCGATTTGGGCATGACC |
| 3783 | catatagggatccatcgtccgacgAAGATCCCCTGATTCCCTTTGT |
| Primers for plasmid pLD3777 (used to make $P_{fur2}$-*gfp*) | |
| 3269 | ccccgggctgcaggaattccGGCACCAGCTACATCCAAC |
| 3270 | ttgtaccgggcccaagcttcCGTGACGCTCCTTTCGTG |
| Primers for plasmid pLD4757 (used to make $P_{glcD}$-*mScarlet*) | |
| 4449 | acgtacgtacactagtTCGAGCAGGTCGTAGAGGGT |
| 4450 | acgtacgtacgaattcGGCGGCTGTCCTTGTTGTG |
| Primers for plasmid pLD4132 (Δ*lldD*) | |
| 3710 | aggcaaattctgttttatcagaccgcttctgcgttctgatAAACCTTTCAAGGCCCTGTT |
| 3711 | ttcgatcagttcgagcaggttCAGGGTGTACTCGGCGTA |
| 3712 | tacgccgagtacaccctgAACCTGCTCGAACTGATCGAA |
| 3713 | ggaattgtgagcggataacaatttcacacaggaaacagctATCAGGTGGGTGAGGATGTC |
| Primers for plasmid pLD3690 (Δ*lldS*) | |
| 2986 | aggcaaattctgttttatcagaccgcttctgcgttctgatCAGGAACGCATCGCGATTCC |

| | |
|---|---|
| 3198 | tcatgcccaaatcgagcagGCTGCCGGTGATCGACAGG |
| 3199 | cctgtcgatcaccggcagcCTGCTCGATTTGGGCATGA |
| 2989 | ggaattgtgagcggataacaatttcacacaggaaacagctCAGCGGCGCTTGATCCATT |
| Primers for plasmid pLD3797 (*lldS* complementation) | |
| 2986 | aggcaaattctgttttatcagaccgcttctgcgttctgatCAGGAACGCATCGCGATTCC |
| 2989 | ggaattgtgagcggataacaatttcacacaggaaacagctCAGCGGCGCTTGATCCATT |
| Primers for plasmid pLD5150 (LldS$^{T107M}$ point mutation) | |
| 4569 | ccaggcaaattctgttttatcagaccgcttctgcgttctgatCGCGGGTTGCTGGCTT |
| 4570 | gcggaatcaggatctgccgggcgAACATCGGCGGCGTGC |
| 4571 | cctgcgcacgccgccgatgttCGCCCGGCAGATCCTGAT |
| 4572 | aattgtgagcggataacaatttcacacaggaaacagctCCGTGGCGATGTCCTCCA |
| Primers for plasmid pLD5173 (LldS$^{T107A}$ point mutation) | |
| 4569 | ccaggcaaattctgttttatcagaccgcttctgcgttctgatCGCGGGTTGCTGGCTT |
| 4566 | cgcggaatcaggatctgccgggcgAAGGCCGGCGGCGTG |
| 4567 | cctgcgcacgccgccggccttCGCCCGGCAGATCCTGAT |
| 4568 | aattgtgagcggataacaatttcacacaggaaacagctGGCCATCTCCTGGCCGAG |
| Primers for plasmid pLD3512 (Δ*lldR*) | |
| 2877 | caggcaaattctgttttatcagaccgcttctgcgttctgatATGGCGCCGATCTTGAAGG |
| 2878 | ggatgtgctggttggacaccCCTCCAGTTGCGCAACGATG |
| 2879 | catcgttgcgcaactggaggGGTGTCCAACCAGCACATCC |
| 2880 | ggaattgtgagcggataacaatttcacacaggaaacagctCAGGAACTCCGGATAGCGCT |
| Primers for plasmid pLD4660 (Δ*fur2*) | |
| 4109 | ggcaaattctgttttatcagaccgcttctgcgttctgatAAGTCCCTTGCCCGCG |
| 4110 | ctgctcgttggccgcgaGGCCGGCCTCCTTCAA |
| 4111 | ttgaaggaggccggcctCGCGGCCAACGAGCAG |
| 4112 | aattgtgagcggataacaatttcacacaggaaacagctTGGCGGCGGCTTCCAG |
| Primers for pLD4600 ($P_{lldA}$*(256)-gfp*) | |
| 4117 | acgtacctcgagGCAGTCCACTCCTTCGGG |

| | |
|---|---|
| 4120 | acgtacactagtGGTGGTGCTGTTCCCGC |
| Primers for pLD4966 ($P_{lldA}$*(221)-gfp*) | |
| 4117 | acgtacctcgagGCAGTCCACTCCTTCGGG |
| 4486 | acgtacactagtACCATTACCACCTGCGCA |
| Primers for pLD4967 ($P_{lldA}$*(188)-gfp*) | |
| 4117 | acgtacctcgagGCAGTCCACTCCTTCGGG |
| 4487 | acgtacactagtCGGCCAGCCAGCGTTT |
| Primers for pLD5029 ($P_{lldA}$*(164)-gfp*) | |
| 4117 | acgtacctcgagGCAGTCCACTCCTTCGGG |
| 4488 | acgtacactagtGGCTGGCGGCGATCT |
| Primers for pLD4599 ($P_{lldA}$*(125)-gfp*) | |
| 4117 | acgtacctcgagGCAGTCCACTCCTTCGGG |
| 4118 | acgtacactagtGGGAATTCCTTCCGGCGG |
| Primers for pLD4598 ($P_{lldA}$*(105)-gfp*) | |
| 4117 | acgtacctcgagGCAGTCCACTCCTTCGGG |
| 4119 | acgtacactagtGACGACTGAAACGGCGGA |
| Primers for pET28a-lldS | |
| LldS-F | aaaacatatgctcaatcccgacgatttc |
| LldS-R | aaaagtcgactcagtcgtccgccgcc |
