## Supplemental file 1 for "The L-lactate dehydrogenases of *Pseudomonas aeruginosa* are conditionally regulated but both contribute to survival during macrophage infection"

**Supplemental File 1**. Pseudomonad L-iLDH profiles. Left: Phylogram representing the relationships between LldD and LldA proteins produced by each of the indicated strains. Center and right: L-iLDH profiles of each of the listed strains. Color shading corresponds to the genomic arrangements shown in the key at the top and in Figure 4A.

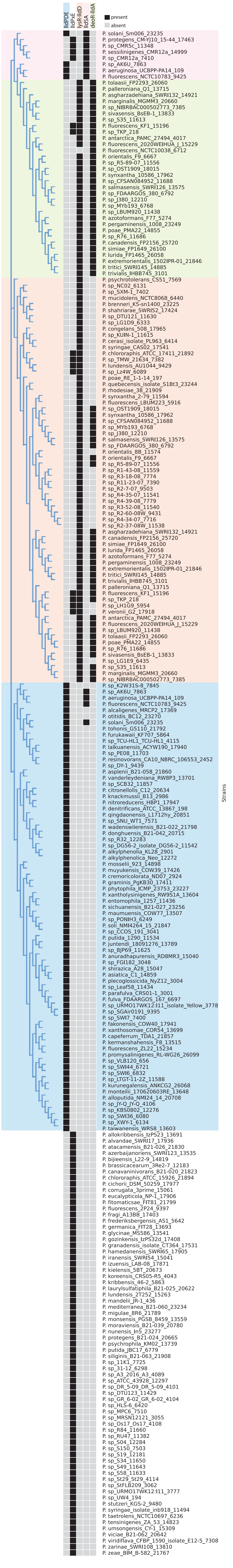
